# A molecular atlas of cell-type specific signatures in the parkinsonian striatum

**DOI:** 10.1101/2025.09.09.675138

**Authors:** Marta Graziano, Ioannis Mantas, Yuvarani Masarapu, Solène Frapard, Raquel Garza, Anita Adami, Shaline Fazal, Annelies Quaegebeur, Roger Barker, Johan Jakobsson, Stefania Giacomello, Konstantinos Meletis

## Abstract

The progressive degeneration of dopaminergic projections to the striatum is a key disease mechanism in Parkinson’s disease (PD). To define the cellular landscape in the parkinsonian striatum, we mapped the cell-type specific transcriptional landscape in early and progressive PD mouse models and in human PD stages. Our analyses revealed substantial transcriptomic changes across both neuronal and glial populations, with astrocytes and oligodendrocytes exhibiting distinct disease-associated gene expression profiles. Notably, progressive dopamine depletion uncovered differential neuronal vulnerability, identifying eccentric striatal projection neurons (SPNs) and Chst9-expressing direct-pathway SPNs as among the most resilient subtypes in both species. This cross-species resource establishes a comprehensive framework for investigating cell-state dynamics in the parkinsonian striatum and uncovers selectively vulnerable and resistant cell types that can inspire new therapeutic strategies.

## INTRODUCTION

Parkinson’s disease (PD) is a neurodegenerative disorder defined in part by basal ganglia (BG) dysfunction initiated through the progressive loss of nigral A9 dopaminergic (DAergic) neurons which extend axonal projections to the striatum^1,2^. As the input nucleus of the BG, the striatum serves as a central hub where DAergic axon terminals broadcast signals related to reward and motor programs^3^. The BG network is traditionally organized into distinct subcircuits based on striatal projection neuron (SPN) types and the dopamine (DA) receptor they express^1^. SPNs expressing DA D1 (Drd1) receptors project directly to BG output nuclei, forming the direct pathway (dSPNs). In contrast, SPNs expressing DA D2 (Drd2) receptors influence BG output indirectly through intermediate BG nuclei, forming the indirect pathway (iSPNs) ^1^. In PD, reduced striatal DA levels are thought to decrease dSPN activity and increase iSPN activity, jointly contributing to some of the major motor features around bradykinesia and rigidity^2,4^. However, owing to the progressive nature of DAergic degeneration in PD, the dynamic changes in SPNs and other cell types in response to gradual DA loss remain unexplored. This is particularly important, as striatal changes may contribute to complications associated with long-term DA-targeted therapies, including L-DOPA-induced dyskinesias^5^.

Molecular profiling using single-cell RNA sequencing and cell-type-specific tracing has been used to map the complex cell types and their connectivity in BG circuits, revealing new cell types embedded within these core pathways^6–11^. Studies have shown that the transcriptional profiles and functional characteristics of SPNs vary along the dorsolateral-ventromedial axis^8,10,11^. Furthermore, the molecular identity and functional roles of SPNs are further shaped by their location within neurochemically distinct compartments, the striosome and matrix^8,10,11^. These findings add new layers to the traditional dSPN-iSPN framework and highlight the need to investigate how DA loss in PD affects transcriptionally distinct cell types and subregions and how these changes contribute to the progression of disease states.

In this study, we aimed to establish a comprehensive cross-species resource on how PD progression impacts cell types and the transcriptional landscape in early and progressive mouse models of dopaminergic denervation (DA depletion) that are widely used as PD models and are based on DA loss rather than α-synuclein pathology, as well as in human PD. To achieve this, we performed spatial transcriptomic (ST) analysis and single-nucleus (sn)RNA sequencing in mild and progressive PD mouse models using different approaches to induce DA degeneration. Specifically, we used a mild toxin model based on low-dose 6-hydroxydopamine (6-OHDA) injections into the medial forebrain bundle, and we used a DA-specific mitochondrial deficiency model (MitoPark mice^12^) at two timepoints of degeneration (10–11 and 15–18 weeks old) to track events during disease progression. We further performed snRNA sequencing from postmortem human striata (putamen) from PD patients across several Braak stages (Braak stage 3 to 6 for Lewy body and neurite pathology^13^) and from age-and sex-matched neurologically healthy donors. In summary, we have established a comprehensive resource that defines the PD-related impact on cell types, and how mild or partial DA degeneration changes the transcriptomic landscape across the striatum.

## RESULTS

### Molecular taxonomy of mouse and human striatal cells in health and parkinsonism

To capture the cell-type-specific transcriptional profile following DA loss at different stages of degeneration, we used PD mouse models with mild or partial DAergic neuron loss (**Figure 1A and 1B**). Specifically, we isolated the caudoputamen (CP) from two PD mouse models with different mechanisms of degeneration: mice with mild 6-OHDA-induced lesions in the medial forebrain bundle and transgenic MitoPark mice at two stages of degeneration (early stage: 10-11 weeks old; middle stage: 15-18 weeks old)^12^ and their respective controls (**Figure 1A and 1B**). We sequenced a total of 160,483 nuclei in total and retained 143,962 nuclei for downstream analysis after quality control filtering (**Figure S1**). On average, we obtained 7,576 nuclei per sample, with an average of 1,992 genes detected per nucleus (**Figure S1**). We identified nine transcriptionally distinct nuclei clusters in the mouse striatum, which we mapped into cell types based on well-established markers (**Figure 1C and 1D**). We were able to identify both major glial and neuronal populations (**Figure 1D**), and the distribution of cell type clusters and the relative proportions of them remained stable across conditions and samples (**Figure S1**). Consistent with previous findings^7,10,11^, dSPNs (*Rgs9+, Drd1+, Tac1+*), iSPNs (*Rgs9+, Drd2+, Adora2a+*) and eccentric SPNs (eSPNs; *Drd1*+, *Col11a1*+, *Sema5b*+) clustered separately (**Figure 1C and 1D**). Additionally, we identified an interneuron cluster (*Npy+, Pvalb+*) and glutamatergic neurons from the deepest layers of the cortex (*Slc17a7+*) (**Figure 1C and 1D**). We also identified several glial populations, such as oligodendrocytes (*Mbp+*), oligodendrocyte precursor cells (OPCs; *Pdgfra+*), astrocytes (*Slc1a3*+, *Aldh1l1+*) and microglia (*Cx3cr1+*) **(Figure 1C and 1D**). We found no condition-specific clusters, as the distribution of cells across clusters was consistent under all conditions (**Figure S1**).

**Figure 1.**
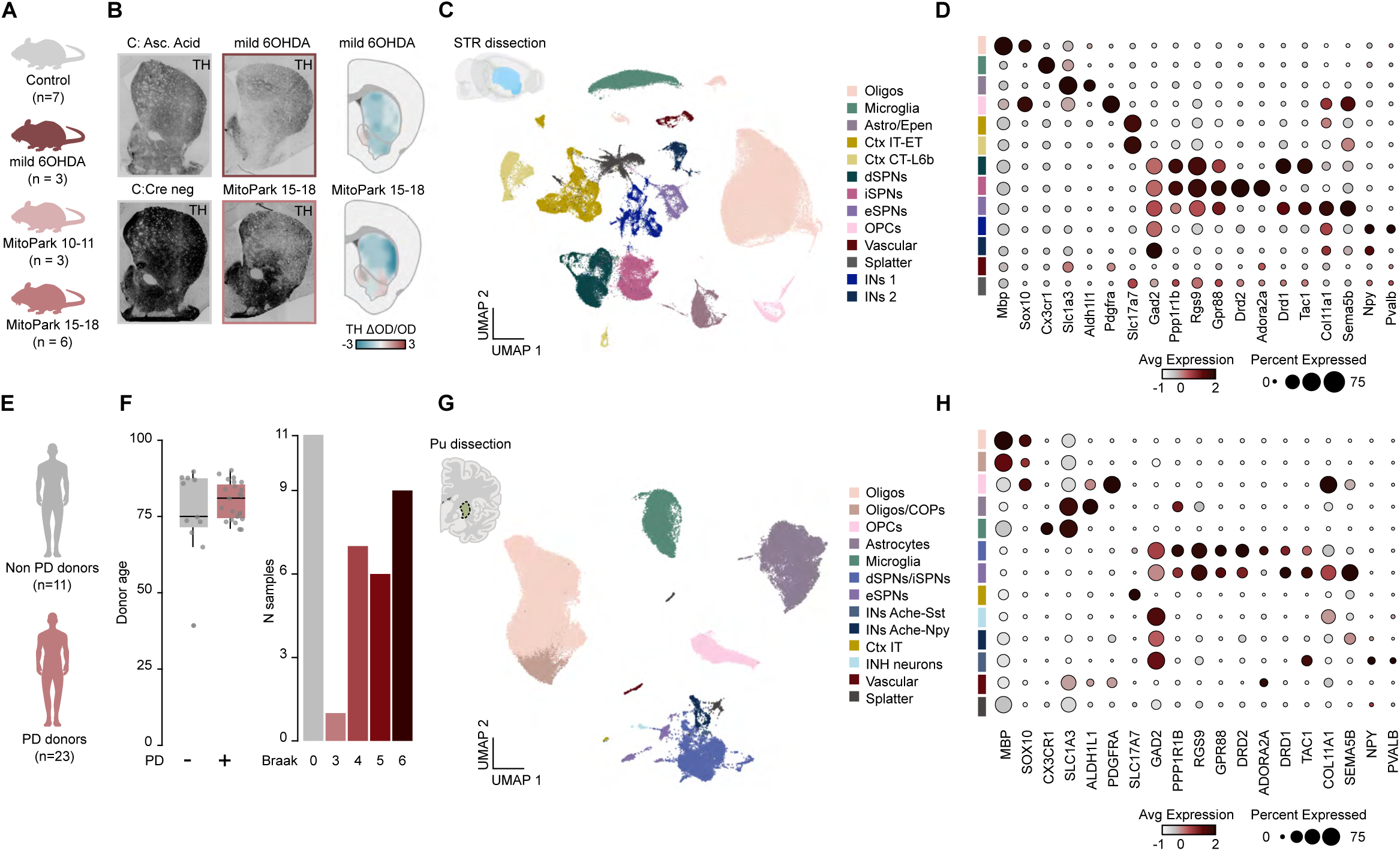
Single-cell atlas of striatal cells from early-stage PD mouse models and PD patients. **A.** Schematic representation of the mouse models used for single-nuclei RNA-seq analysis and respective controls. **B.** Representative images and heatmap showing the intensity of TH loss in the striata of control, mild 6OHDA, and MitoPark 15-18 mice. (n=3 for each group). **C.** UMAP visualization of cell types in the striatal mouse dataset with cluster annotations. **D.** Dot plot showing the cell-type enriched markers in the mouse PD dataset (color-coded as in **C**). **E.** Schematic of the samples collected for the analysis of the collected post-mortem putamen samples from PD donors and non-PD donors used for single-nuclei RNA-seq analysis. **F.** Boxplot showing the distribution of age in the samples, and bar plot showing the number of samples split by Braak stages. **G.** UMAP visualization of cell types in the human putamen dataset with cluster annotations. **H.** Dot plot showing the cell-type enriched markers in the human PD dataset (color-coded as in **D**).

We further isolated nuclei from the putamen (Pu) of PD patients (23) and control subjects (11) and analyzed the transcriptional profiles of the cell types (**Figure 1E**). Control non-PD samples were age matched (**Figure 1F**), and the samples from PD patients included those with Braak stage pathology ranging from 3 to 6 (**Figure 1F, S3**). In this analysis, we isolated 75,875 nuclei and analyzed 75,530 nuclei after filtering (**Figure S2**), with an average of 2,172 nuclei per sample and 2,280 genes per nucleus. We identified seven transcriptionally distinct cell populations based on markers expression (**Figure 1G and 1H**). Specifically, we identified neuronal populations, including SPNs (*RGS9*+, *PPP1R1B*+, *GPR88*+) and interneurons (*SST*+); however, this analysis did not result in clear separation between dSPNs and iSPNs (**Figure 1G and 1H**). For the glial subtypes, we identified oligodendrocytes (*MPB*+), astrocytes (*SLC1A3*+, *ALDH1L1*+), microglia (*CX3CR1*+), and OPCs (*PDGFRA*+) (**Figure 1G and 1H**). The distribution of clusters and proportions of cell types were consistent across all samples, indicating that these patterns are stable and unaffected by variables such as age, sex, or Braak stage (**Figure S3**).

### Transcriptomic effects on neuronal and glial cell types across PD mouse models with partial DAergic neuron depletion

To evaluate the impact of DAergic neuron loss, we performed differential expression (DE) analysis on the entire dataset for each PD mouse model compared to its corresponding control (**Figure 2A-2C**). In the subsequent analysis, we excluded cells identified as the “Splatter” cluster, as they likely represent artifacts. In early-stage MitoPark animals, the DE analysis revealed a smaller number of DE genes (1059 genes; **Figure S4**). Both middle-stage MitoPark mice and mildly lesioned 6-OHDA mice presented a substantial increase in DE genes (MitoPark: 1801 genes, 6-OHDA: 1552 genes; **Figure 2A and 2B**). DE gene analysis across all clusters revealed that MitoPark mice exhibited a relatively balanced expression of upregulated and downregulated genes with a log fold change greater than 0.2 (**Figure 2A and 2B**). In contrast, mice in the mild 6-OHDA model predominantly displayed downregulated genes (**Figure 2A and 2B**). Even though early-stage MitoPark samples displayed low count of DE genes, analysis of the normalized DE genes counts per cluster revealed that oligodendrocytes were the main cluster affected (**Figure S4**). Similar analysis in the neuronal clusters of the 6-OHDA model revealed that exhibited a higher magnitude of change and eSPNs were the least affected neuronal populations (**Figure 2C**). A similar analysis of the neuronal clusters of middle-stage MitoPark mice revealed a uniform impact across most cell populations, with the eSPN cluster remaining the least affected, which was consistent with the findings of the 6-OHDA model analysis (**Figure 2C**). Analysis of normalized DE gene counts in glial clusters revealed oligodendrocytes demonstrated the highest degree of transcriptional variation across both PD models (**Figure 2C**). Compared with that in the 6-OHDA mouse model, the astrocyte/ependymal cluster was predominantly affected in the middle-stage MitoPark model (**Figure 2C**). Moreover, compared with the other clusters, the oligodendrocyte cluster presented the greatest fold changes in DE genes (**Figure 2D and 2E**). Given the observed vulnerability of dSPNs and oligodendrocytes in both PD models, we sought to identify the genes significantly affected in these cell types across both models. This analysis revealed that the most significantly affected genes, based on fold change, were the downregulated ones (**Figure 2F**). Notably, several of these genes, including *Rgs20, Homer1, Prkcb*, and *Dusp14*, are associated with G-protein-coupled receptor (GPCR) signaling. In oligodendrocytes, DE gene analysis in both models revealed that *Apoe* and *Apod*, amyloid-binding-related apolipoproteins, were among the genes most significantly upregulated according to fold change between the models (**Figure 2G**). We next performed pseudotemporal^14^ ordering and trajectory inference analysis^15^ of transcriptional dynamics in clusters derived from the MitoPark samples (including both early and middle stage) to capture molecular changes during progressive DA loss, pseudotime and trajectory inference have traditionally been used to model developmental changes, but more recent studies have applied these methods to investigate disease progression^16,17^. Pseudotime measures the progression of cells along a trajectory in a dimensionally reduced space, but it does not inherently account for gene expression variability or changes. In our analysis, pseudotime reconstruction was applied to the combined dataset of neuronal and glial populations, which differ markedly in transcriptional profiles and developmental origins; however, our aim was to capture broad, cross-cell-type trajectories of transcriptional change.

**Figure 2.**
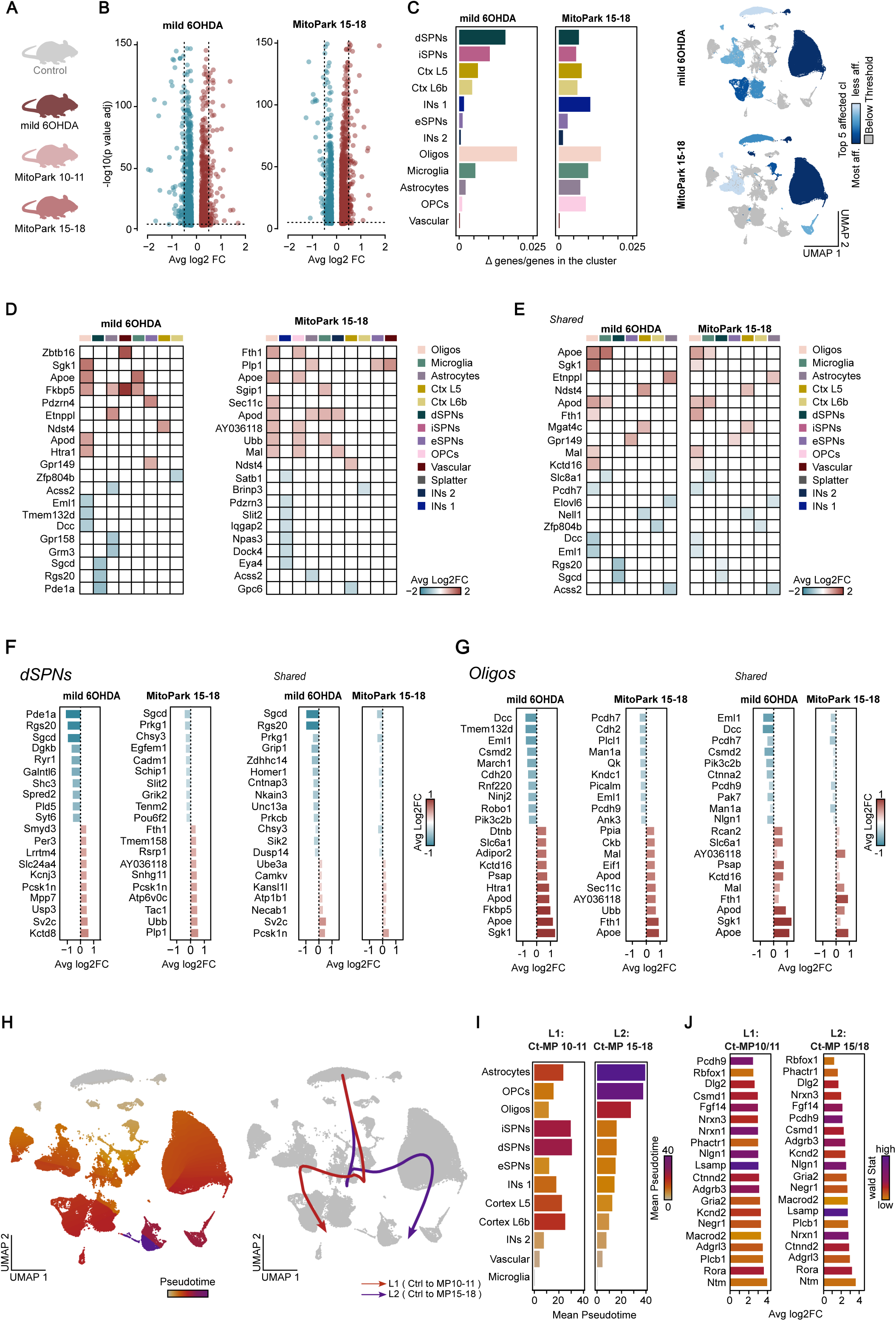
Vulnerability of D1+ SPNs and oligodendrocytes in early-stage PD mouse models. **A.** Schematic of mouse models and controls used for DE analysis of snRNA sequencing. **B.** Volcano plots of differentially expressed genes in the mild 6-OHDA model and MitoPark model, (15–18 weeks; p < 0.0001, log₂FC > 0.2 or < −0.2). **C.** Bar plots of significantly changed genes over total genes per cluster in the mouse models, with neuronal and glial identities indicated (left); UMAP visualization of the top five affected clusters (right). **D-E.** Heat maps of the top 20 dysregulated genes in the mild 6-OHDA and MitoPark models (D), and of shared dysregulated genes (E), grouped by cluster and colored by average log₂FC. **F.** Bar plots of average log₂FC for the top 20 dysregulated genes in dSPNs per mouse model, and for shared genes across models. **G.** Bar plots of average log₂FC for the top 20 dysregulated genes in oligodendrocytes per mouse model, and for shared genes across models **H.** UMAPs colored by pseudotime progression (left) and inferred trajectories (right), with curves indicating lineage paths. **I.** Mean pseudotime values per cluster across lineages, ordered and color coded by trajectory. **J.** Bar plots of average log₂FC for the top 20 pseudotime-associated genes per lineage, colored by correlation with pseudotime (wald stat).

To define the trajectory, we used control samples as the starting point and middle-stage MitoPark samples as the endpoint. We reconstructed the transcriptional response trajectories of striatal cells along a pseudotime axis, revealing two distinct lineages corresponding to early (lineage 1) and middle (lineage 2) stages of MitoPark (**Figure 2H-2J**). Both lineages originated from the microglia cluster, with lineage 1 terminating in the SPN clusters and lineage 2 terminating in the astrocyte/ependymal cluster (**Figure 2H-2J**). These findings suggest that dSPNs undergo significant transcriptional changes early in DA denervation, whereas distinct glial responses emerge as the disease progresses.

### Transcriptomic effects on SPN subtypes across PD mouse models with partial DA neuron depletion

To investigate in detail the transcriptional changes in the distinct subtypes of SPNs, we proceeded to subclustering the dSPN, iSPN and eSPN populations (**Figure 3A**). This analysis was successful in revealing the main parcellation of SPNs, in accordance with previous studies (**Figure 3A and 3B**)^7,10,11^. Specifically, we observed a clear separation in the UMAP landscape of dSPNs and iSPNs that belong to the matrix (*Id4+, Epha4+*) and patch *(Sema5b+, Lypd1+, Kremen+*) compartments (**Figure 3A and 3B**). Consistent with previous studies, we identified clusters defining the dorsolateral striatum (DLS; *Gpr155* enriched) and dorsomedial striatum (DMS; *Crym* enriched) matrix for both dSPNs and iSPNs, as well as a patch-iSPN subtype characterized by strong *Htr7* expression (**Figure 3A and 3B**)^7,10^. Moreover, we were able to distinguish the clusters that belonged to the ventral striatum (VS; *Dlk1, Crym, Syt10* and *Gabra5* enriched), such as the dSPN-VS, iSPNs-VS and striatum-like amygdala nuclei (sAMY) clusters. The islands of Calleja formed a distinct cluster with a unique transcriptional profile compared to the VS with marked expression of *Drd3* and *Col11a1*, a finding consistent with previous reports^10^. We further identified two transcriptionally unique clusters of SPNs that were located away from the remaining SPNs, and the gene expression profiles of these unique SPN clusters resembled those of eSPNs (*Col11a1+, Otof+, Pcdh8+, Sema5b+*)^11^ and the recently identified Chst9+ SPNs. The eSPN gene expression profile also matched the expression profile of the *Drd1*-*Drd2* hybrid population reported in other studies (**Figure 3A and 3B**)^7,10^. Additionally, we identified a cluster expressing both glial (*Mbp*+) and SPN-specific markers, which we determined was a technical artifact and excluded from further analysis. DE analysis of the entire SPN dataset revealed a moderate number of DE genes in the mild 6-OHDA model (457; **Figure 3C**; **Figure S5**), with a predominance of downregulated genes. In contrast, the middle-stage MitoPark model exhibited a less pronounced transcriptional response (329; **Figure 3C**; **Figure S5**), characterized by a slight tendency toward upregulation. Based on the number of DE genes, we observed a dorsoventral gradient, with clusters representing the dorsal striatum (DS) being more affected than those representing the VS (**Figure 3D**). Notably, the matrix DMS-dSPN cluster is the one that showed the largest changes in transcription (**Figure 3D**). Conversely, ventral striatal clusters, along with Htr7+ patch-iSPNs, eSPNs and Chst9+ SPNs, demonstrated resilience to DA neuron loss (**Figure 3D**). An examination of the shared genes with the greatest fold changes across both models revealed that these genes were predominantly identified within the matrix DMS-dSPNs (**Figure 3E and 3F**). Analysis of the DE genes in the matrix DMS-dSPNs of both the mild 6-OHDA and middle-stage MitoPark models revealed several genes related to GPCR signaling, with key genes being *Rgs20, Homer1* and *Prkcb*. Analysis of DE genes in the matrix DLS-dSPNs of both the mild 6-OHDA and middle-stage MitoPark models revealed several genes with downregulated expression related to synaptic integrity (*Cntnap3, Nrxn3, Unc13a*) and neuronal activity (*Nr4a1, Celf2*) (**Figure 3G and 3H**). To investigate the differential vulnerability of SPN subtypes to DA neuron loss, we performed pseudotemporal analysis on clusters from control samples and the two stages of the MitoPark model. Through this analysis, we identified two distinct lineages corresponding to the early (lineage 1) and middle (lineage 2) stages of the MitoPark model (**Figure 3I-3J**). Both lineages originated from the eSPN cluster, with lineage 1 terminating in the matrix DLS-dSPN cluster and lineage 2 terminating in the matrix DLS-iSPN cluster (**Figure 3I-3J**). These results suggest that matrix DLS-iSPNs could be affected at earlier stages of DA depletion (early-stage MitoPark), whereas matrix DMS and DLS-dSPNs are highly affected at a more advanced stage of DA loss (middle-stage MitoPark). Accordingly, we observed that numerous trajectory-influencing genes across sample groups in both lineages were enriched within the eSPN (e.g., *Adarb2, Sema5b*) cluster (**Figure 3K**). Collectively, these findings suggest that dorsal striatal matrix SPN type shows early vulnerability during partial DAergic degeneration, whereas eSPNs represent the most resilient SPN type.

**Figure 3.**
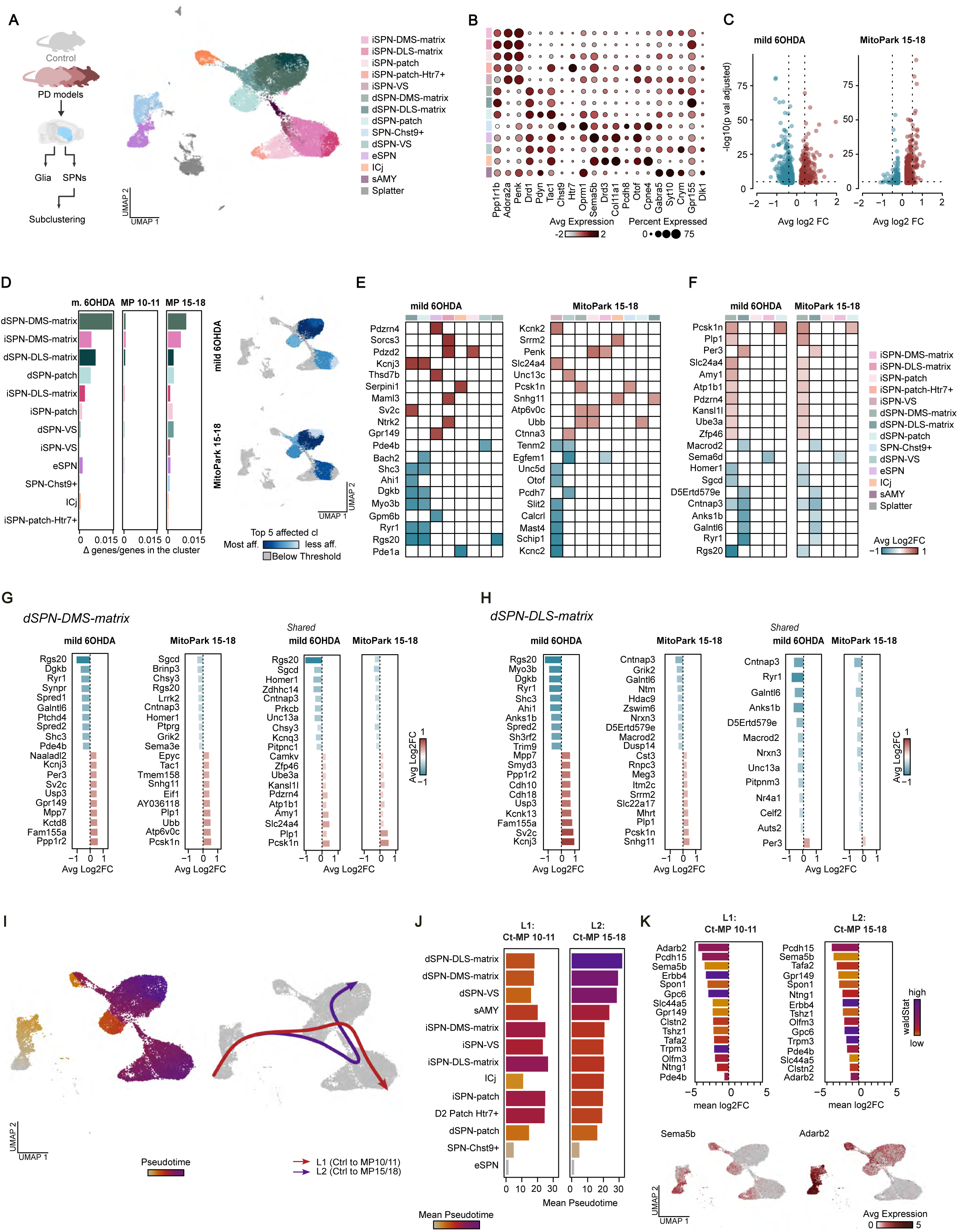
Differential PD vulnerability of patch and matrix striatal compartments. **A.** Schematic of the experimental and analytical workflow (left), with subclustering of SPNs and UMAP projection color coded by cluster annotation (right). **B.** Dot plot showing cell-type enriched markers in the mouse SPNs subset. **C.** Volcano plots of differentially expressed genes in SPN subclusters from the mild 6-OHDA model and MitoPark model (15–18 weeks; p < 0.0001, log₂FC > 0.2 or < −0.2). **D.** Bar plots of significantly changed genes over total genes per cluster in both models, grouped by neuronal and glial identity; UMAP visualization of the top five affected clusters (right). **E-F**. Heat maps of the top 20 dysregulated genes in SPN subclusters in mild 6-OHDA and MitoPark (E), and of shared dysregulated genes (F), grouped by cluster and colored by average log₂FC. **G.** Bar plots of average log₂FC for the top 20 dysregulated genes in DMS-matrix dSPNs per model, and for shared genes across models. **H.** Bar plots of average log₂FC for the top 20 dysregulated genes in DLS-matrix dSPNs per model, and for shared genes across models. **I.** UMAPs colored by pseudotime progression (left) and inferred trajectories (right), with curves indicating lineage paths. **J.** Mean pseudotime values per cluster across lineages, ordered and color coded by trajectory. **K.** Bar plots of average log₂FC for the top 20 pseudotime-associated genes per lineage, colored by correlation with pseudotime (wald stat); example genes shown on UMAPs.

### Spatial transcriptomic map of a mild DA degeneration model

To explore whether the spatial distribution of transcriptomic changes depends on regional differences in DAergic neuron depletion, we performed spatial transcriptomics (ST) analysis in the mild 6-OHDA lesion model and respective controls (3 mild 6-OHDA, 3 controls; **Figure 4A**). We selected four coronal sections spanning the anteroposterior axis of the striatum (two precommissural and two postcommissural sections) to comprehensively sample the striatal structure (**Figure 4A**). We identified 32 distinct spatial clusters, demonstrating clear separation between cortical (*Slc17a7*+) and subcortical areas (**Figure 4A-4C**; **Figure S6**). Within the striatum, clusters segregated based on spatial markers rather than dSPN or iSPN identity, as *Drd1* and *Drd2* expression was intermingled across clusters. Specific clusters were associated with distinct regions, including the DMS (*Crym*+, clusters 8 and 5), DLS (*Gpr155*+, cluster 4), and VS (*Crym*+, *Dlk1*+, *Col6a1*+, clusters 6 and 24). Additionally, we identified a cluster defining white matter in the corpus callosum (*Mbp*+, cluster 7; **Figure 4A-4C**; **Figure S6**). Then, we aimed to identify which of the spatial clusters are more affected by the mild 6-OHDA lesion. Analyzing consecutive sections to define the loss of DA innervation using TH staining showed an overall decrease in the 6-OHDA-treated striata compared with the control striata, with regional differences showing greater intensity in the lateral striatum and lower intensity in the medial striatum (**Figure 4D**). DE analysis across spatial clusters revealed that 203 DE genes, comprising 121 downregulated and 83 upregulated genes (adjusted p < 0.0001), were distributed across 27 ST clusters. Among the extrastriatal regions, the claustral complex (cluster 16) and cortical layer 5a (cluster 10) presented the greatest number of DE genes (**Figure 4D**). Within the striatum, clusters corresponding to the DMS (clusters 5, 8 and 12) presented the greatest transcriptional differences in DE genes, which was consistent with the snRNA-seq findings in SPNs. DE gene changes in the DS were negatively correlated with tyrosine hydroxylase (TH) immunoreactivity intensity, which was weaker in the DMS than in other striatal regions (**Figure 4D**). Notably, a marked decrease in TH immunoreactivity was observed in the VS in this PD model, but this reduction did not correlate with the number of DE genes in the region (**Figure 4D**). Detailed examination of the genes driving these effects revealed that the mitochondrial function-associated gene *Lars2*^18^ was significantly upregulated, with pronounced expression in layer 5a and the claustrum (**Figure 4E**). In contrast, dorsal striatal clusters exhibited marked downregulation of genes linked to GPCR signaling pathways, including *Nr4a1* and *Egr1* (**Figure 4E**). These findings suggest that the cell-type specific vulnerability in stratum is dictated by the spatial distribution of cell types in striatum in combination with decreased DA levels.

**Figure 4.**
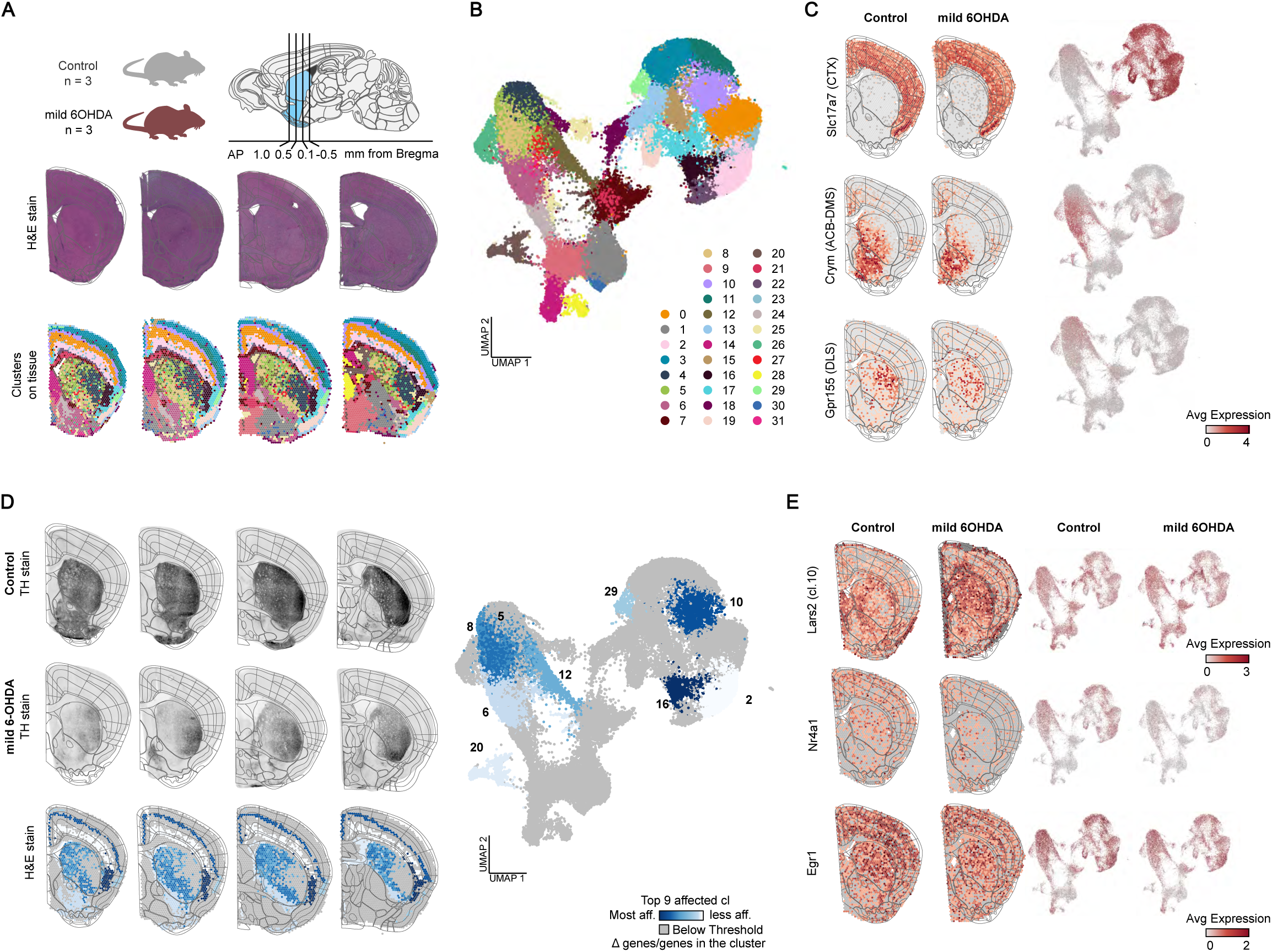
Spatial transcriptomics defines molecular changes in striatal subregions in a mild DA degeneration model. **A.** Schematic of the experimental workflow, showing mouse models, brain sections (Allen Brain Atlas reference), H&E staining, and spot overlays. **B.** Visualization of UMAP projection of spatial spots based on integrated data. **C.** Average expression of cell type markers defining spatial domains, plotted on representative tissue sections (left) and UMAPs (right). **D.** Tyrosine hydroxylase (TH) loss in adjacent sections (as in Fig. 1); bottom row shows spots for the top nine affected clusters, color coded by normalized Δ genes/total genes, and corresponding UMAP (p < 0.0001). **E.** Average expression of representative differentially expressed genes in the most affected clusters, shown as spots overlaid on tissue (left) and UMAPs (right), split by condition (control vs. mild 6-OHDA).

### Transcriptome profile of neuronal and glial cell types in the human PD Putamen

Given that PD mouse models do not fully capture the complexity of the human disease, we generated and analyzed a large snRNA dataset from the human putamen, including PD and non-PD cases (**Figure 5A**). When performing differential expression analysis and applying the same adjusted p-value and log₂ fold-change thresholds as in the mouse dataset, we detected 2,681 dysregulated genes **(Figure 5B**). Among the different cell types, SPNs exhibited the greatest transcriptional changes in PD samples, whereas astrocytes and microglia showed the largest effects among glial populations (**Figure 5C**). Notably, the 20 genes with the most significant fold-change alterations were not confined to the SPN or astrocyte clusters (**Figure 5D**). Given the observed vulnerability of SPNs and astrocytes, we examined the genes significantly affected in these cell types, revealing their involvement in diverse biological pathways and functions (**Figure 5E**). Using the non-PD samples as the starting point and the PD samples with advanced Lewy body and neurite pathology (Braak stage 6) as the endpoint, we reconstructed the transcriptional response trajectories of striatal cells along a pseudotime axis, with the same rationale used for the mouse dataset. As Braak staging primarily reflects α-synuclein pathology, it may not directly correspond to the extent of dopaminergic neuron loss; in this context, we use it as a general framework for ordering disease progression. This analysis identified two distinct lineages: one corresponding to Braak stage 5 (lineage 1) and the other to Braak stage 6 (lineage 2; **Figure 5F and 5G**). Both lineages originated from the SPN cluster and terminated in the astrocyte cluster (**Figure 5F and 5G**). However, their trajectories differed slightly, with lineage 1 reaching the astrocyte cluster via microglia, while lineage 2 passed through OPCs before reaching the astrocyte cluster (**Figure 5F and 5G**). Overall, these findings suggest that SPNs show the first transcriptional changes in the human PD putamen, while astrocytes exhibit significant gene expression changes at more advanced points in disease progression.

**Figure 5.**
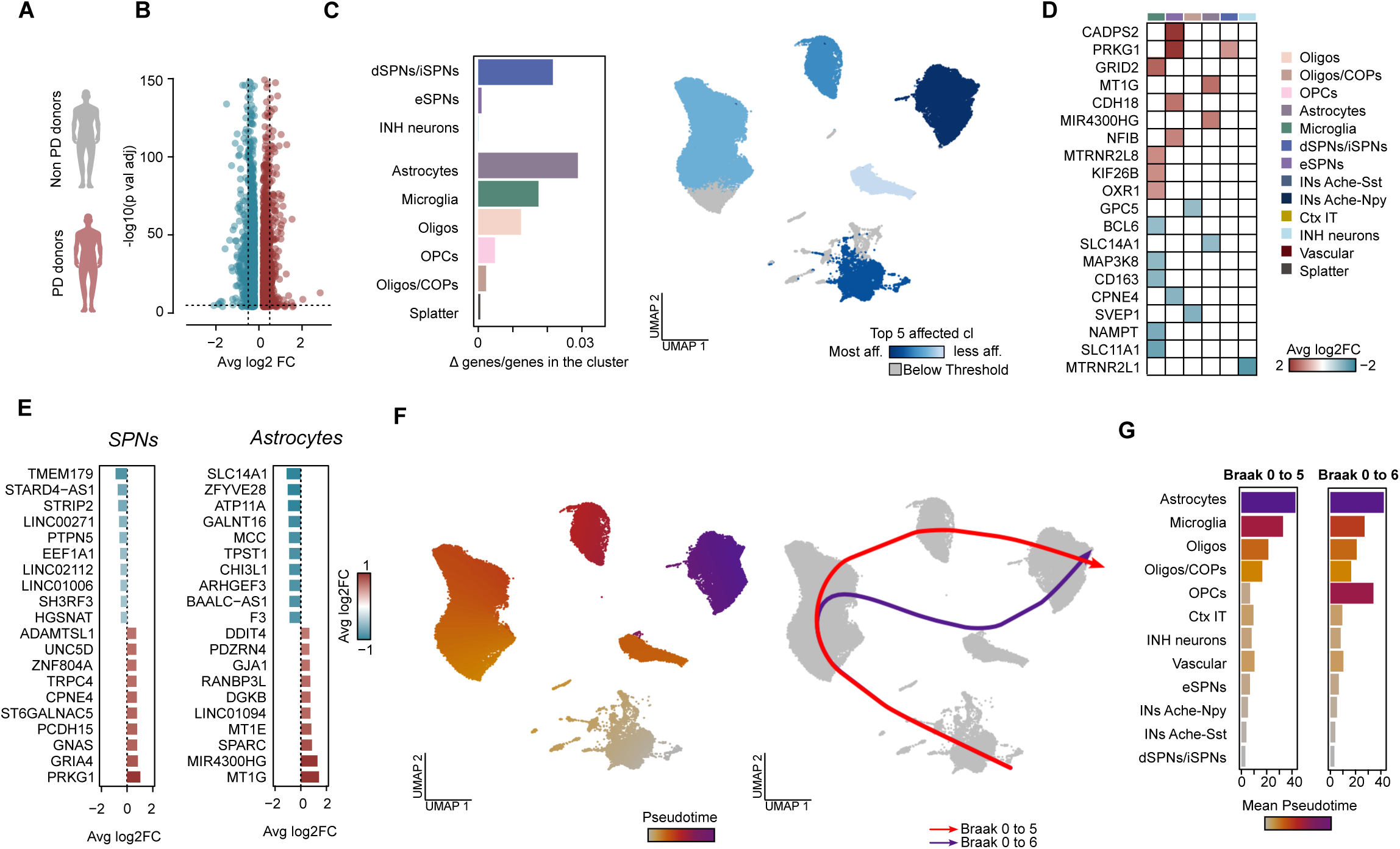
Molecular profile of striatal cell types in human PD. **A.** Schematic of post-mortem putamen samples from PD and non-PD donors used for single-nucleus RNA-seq. **B.** Volcano plot of differentially expressed genes in PD versus controls (p < 0.0001, log₂FC > 0.2 or < −0.2). **C.** Bar plots of significantly changed genes over total genes per cluster normalized Δ genes/total gene, grouped by neuronal and glial identity (left); UMAP visualization of the top five affected clusters (right). **D.** Heat map of the top 20 dysregulated genes and their cluster assignments, colored by average log₂FC. **E.** Bar plots of average log₂FC for the top 20 dysregulated genes in the two most affected clusters: SPNs (neuronal; left) and astrocytes (glial; right). **F.** UMAPs colored by pseudotime progression (left) and inferred trajectories (right), with curves indicating lineage paths. **G.** Mean pseudotime values per cluster across lineages, ordered and color coded by trajectory

### Transcriptome profile of SPN subtypes across Braak stages in the human PD putamen

To investigate the transcriptional changes in distinct SPN subtypes in greater detail, we performed subclustering of the SPN and eSPN populations (**Figure 6A**). This analysis successfully revealed the primary parcellation of SPNs, integrating insights from previous snRNA-seq studies on macaque and human striata^6,8^, as well as murine snRNA-seq datasets (**Figures 6B**)^7,10^. Specifically, we observed a clear separation in the UMAP landscape between dSPNs *(DRD1*+, *TAC1*+) and iSPNs (*DRD2*+, *ADORA2A*+), which were further distinguished by their matrix (*KCNIP1* low) and patch (*KCNIP1* high) compartments (**Figures 6B**). Additionally, we identified a human equivalent of the murine *Htr7*+ patch-iSPN subtype (*DRD2*+, *HTR7*+, *KCNIP1*+; **Figures 6B**). Consistent with previous study findings, we identified two transcriptionally distinct SPN clusters that segregated clearly from the main SPN populations based on their unique gene expression profiles. These clusters displayed characteristics of eSPNs (also known as D1/D2-Hybrid) previously described in macaque and human striata (*COL11A1+, SEMA5B+, RXFP1+, CASZ1+)*^6,8^. The second cluster resembled the gene expression profile of D1-NUDAP cells described in macaque striata (*STXBP6*+, *CHST9*+)^8^, likely representing the primate equivalent of murine *Chst9*+ SPNs (**Figure 6B**). We also identified an SPN cluster that did not align perfectly with previously described populations (SPN-mixed cluster). Finally, we identified a few clusters that did not fully conform to the SPN-specific profile, which we classified as likely technical artifacts. DE analysis of the entire SPN dataset revealed a substantial number of DE genes in the PD samples (2,679 for full dataset; **Figure 6C**, 1,819 and 2,571 for respectively comparing non-PD donors with Braak 4 and Braak 6; **Figure S7**). Among the SPN subtypes, matrix iSPNs demonstrated the highest degree of transcriptional variation, followed by matrix dSPNs, based on the number of DE genes (**Figure 6D**). Interestingly, we observed that eSPNs and *CHST9*+ dSPNs are resistant to PD changes as they exhibited a strikingly smaller number of DE genes (**Figure 6D**). The majority of the 20 genes most significantly affected in PD, based on fold change, were predominantly localized in the ICj and SPN mixed clusters (**Figure 6E**). Further analysis revealed that matrix iSPNs and dSPNs shared several of the highest upregulated (*PRKG1, MGAT4C, SLC35F1, ADAMTSL1, PWRN4, ST6GALNAC5*) and downregulated (*TMEM179, STRIP2, STARD4-AS1, HGSNAT, EEF1A1*) genes by fold change (**Figure 6F**). We next examined the common DE genes significantly affected in both the mouse PD models and PD donors (**Figure 6G**). We assumed that the mild 6-OHDA and middle-stage MitoPark models resemble closely Braak stage 4. Therefore, we compared the top 20 shared DE genes by fold change in Braak stage 4 samples with those in mild 6-OHDA and middle-stage MitoPark samples separately. In the Braak stage 4 matrix dSPNs, we observed overlap among several downregulated DE genes found in mouse dSPN clusters from the 6-OHDA model, including *SPRED1*, *PDE1B* and *SH3RF2* (**Figure 6G**). Likewise, several downregulated DE genes from Braak stage 4 matrix dSPNs overlapped with DE genes found in mouse dSPNs from the middle-stage MitoPark model, such as *TMEM179*, *TARID* and *NRG1* (**Figure 6G**). Overall, these findings suggest some overlapping transcriptional changes in the striata of PD donors at moderate pathological stages and mouse PD models with partial DA neuron depletion, which aligns with Braak stage 4. To investigate the differential vulnerability of SPN subtypes across varying severities of PD pathology, we performed a pseudotemporal analysis of these clusters. We defined non-PD samples as the starting point and PD samples with Braak stage 6 as the endpoint to reconstruct transcriptional response trajectories along a pseudotime axis. This analysis revealed a lineage originating from the matrix dSPN cluster and extending toward the eSPN/CHST9+ SPN clusters (**Figure 6H, 6I**). These findings suggest that matrix dSPNs exhibit significant transcriptional changes at earlier pseudotime values, whereas eSPNs and CHST9+ SPNs undergo distinct alterations at more advanced pseudotime values, potentially reflecting later-stage vulnerability. Interestingly, genes strongly correlated with this trajectory (e.g., *CNTN5, LUZP2, FOXP2*) were enriched in the eSPN and CHST9+ SPN clusters (**Figure 6J**), further supporting their involvement in later transcriptional changes. Collectively, these results suggest that in humans, dorsal striatal matrix SPNs might be particularly susceptible to PD, while eSPNs and CHST9+ SPNs may show greater transcriptional shifts at advanced pseudotime values.

**Figure 6.**
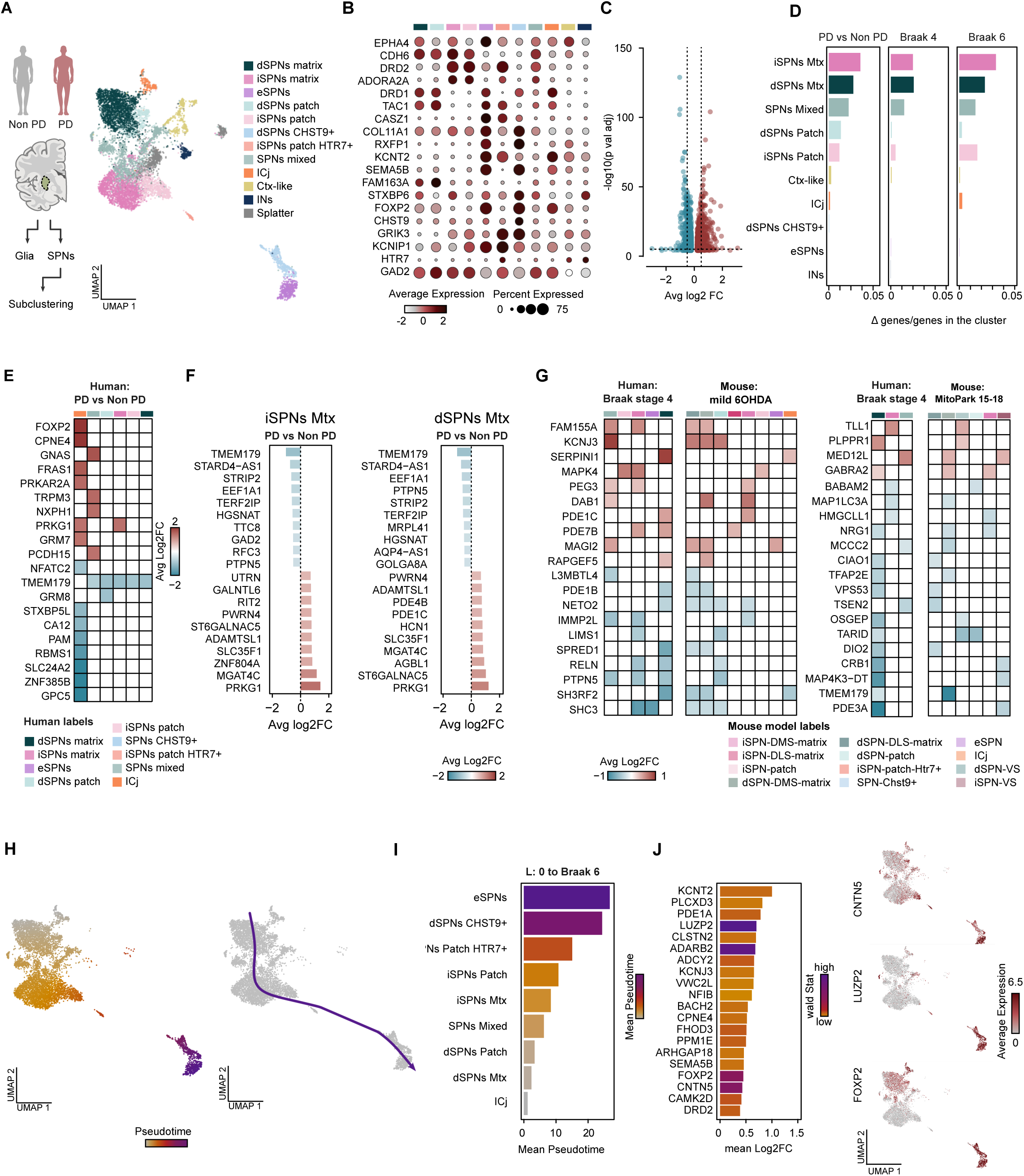
Transcriptome profile of SPN subtypes across Braak stages in the human PD. **A.** Schematic of the experimental and analytical workflow with subclustering of SPNs from human single-nucleus RNA-seq (left); UMAP projection color coded by cluster annotation (right). **B.** Dot plots showing expression of selected marker genes across SPNs clusters, used for cluster annotation. **C.** Volcano plots of differentially expressed genes in SPNs subclusters from PD versus non-PD donors (p < 0.0001, log₂FC > 0.2 or < −0.2). **D.** Bar plots of significantly changed genes over total genes in human SPNs, comparing PD versus non-PD donors and stratified by Braak stage (4 and 6). **E.** Heat maps of the top 20 dysregulated genes in the SPN subset, colored by average log₂FC **F.** Bar plots of average log₂FC for the top 20 dysregulated genes in iSPNs (left) and dSPNs (right). **G.** Heat maps of commonly affected genes in SPN subclusters from human Braak stage 4 and mild 6-OHDA model (right) or MitoPark model (15-18 weeks old; left), grouped by cluster and colored by average log₂FC. **H.** Mean pseudotime values per cluster across lineages, ordered and color coded by trajectory. **I.** Bar plot of the mean pseudotime plotted for each cluster in each lineage (trajectory), color coded and ordered by mean pseudotime. **J.** Bar plots of average log₂FC for the top 20 pseudotime-associated genes per lineage, colored by correlation with pseudotime (wald stat; left); example genes shown on UMAPs (right).

## DISCUSSION

In this study, we have established a comprehensive molecular map of healthy and pathological cell states in the striatum from PD mouse models and PD patients. This resource is based on a comprehensive and unbiased sampling of mouse striatal cells (more than 140,000 cells) and neurons (21,248 SPNs), offering valuable insights into cell-type-specific responses to disease progression. We employed two distinct PD mouse models with different neurodegeneration mechanisms: the 6-OHDA model (neurotoxin-based^8^) and a dopamine-specific mitochondrial deficit model (MitoPark mice^12^). The progressive nature of degeneration in the MitoPark model enabled us to examine two stages of neurodegeneration, capturing cell-specific changes throughout the neurodegenerative trajectory. This snRNA-seq dataset is complemented by an ST dataset of striata from the partial 6-OHDA model, offering a valuable resource for investigating changes affecting the spatially segregated subregions of the striatum. We further provide a large dataset on healthy and PD-affected Putamen, integrating over 70,000 cells from healthy and PD donors, approximately 7,184 of which were identified as putative SPNs. Importantly, our human data includes samples from PD donors across different Braak pathology stages, offering critical insights into cell-type-specific responses to PD progression. This combined resource has the potential to inform the development of new therapeutic cell-type specific interventions targeting PD.

Our study extends previous transcriptomic analyses in PD, which have predominantly focused on acute lesion models. For example, the acute neurotoxin-induced lesion models (6-OHDA or 1-methyl-4-phenyl-1,2,3,6-tetrahydropyridine (MPTP) have been studied; however, these models cannot recapitulate the progressive nature of the disease or the distinct pathophysiological mechanisms underlying neurodegeneration^8,19–22^. In this study, we focused on a mitochondrial deficiency PD model (MitoPark mice) with a slow DAergic neuron degeneration pattern resembling human PD progression^12^. To capture the progression of the striatal transcriptional state, we compared early-and mid-stage degeneration stages and developed a comprehensive high-resolution trajectory map of the cell clusters affected at different stages of degeneration. Importantly, we collected and analyzed human putamen samples from various Braak stages, which represent different degeneration stages^13^. This approach enables a direct comparison between the dynamic transcriptional and cell state effects in mouse PD models and corresponding human PD stages, thereby providing critical insights into the validity and relevance of these models for understanding PD pathophysiology. It also enables us to understand in PD samples what is driven by the disease process rather than DAergic therapy, given the patients will be on a range of DAergic agents for their condition.

SPNs are the primary targets of DA signals; however, our findings demonstrate that there are significant transcriptomic changes in glial populations within the striatum and surrounding tissue in mouse as well as human PD tissue. Our analysis revealed that, compared with microglia, there were significant alterations in astrocytes within both middle-stage MitoPark model mice and human PD striata. Moreover, in both middle-stage MitoPark and human PD striata, astrocyte clusters were primarily affected during the later stages of DAergic degeneration. Importantly, our findings align with studies reporting reactive astrocytes in caudate PD patients^23^. Emerging evidence suggests that disruptions in astrocyte biology contribute to DAergic neuron degeneration in PD. Astrocytes play a neuroprotective role by releasing neurotrophic factors, producing antioxidants, and clearing neuronal waste products, including damaged mitochondria^24^. The latter function is particularly relevant to our study, as the MitoPark model is based on mitochondrial deficits restricted to DAergic neurons^12^. This finding suggests that striatal astrocytes may facilitate the clearance of dysfunctional mitochondria released from dying DA axon terminals in both MitoPark models and PD patients. Moreover, the observed response of striatal astrocytes to DA loss may also involve their role in DA metabolism^25^. Astrocytes are known to be activated by DA and express enzymes involved in its catabolism, such as monoamine oxidase B^26,27^. These characteristics suggest that astrocytes may represent a particularly vulnerable glial cell type in both the MitoPark model and human PD.

However, our findings from glial populations in the mild 6-OHDA PD model differ from human data, as oligodendrocytes, rather than astrocytes, emerged as the most prominently affected glial cell type. Notably, oligodendrocytes were also identified among the most affected glial populations in middle-stage MitoPark models. Recent evidence implicates oligodendrocytes in the substantia nigra during the early stages of PD, which is supported by associations with PD risk genes and transcriptional alterations^28,29^. Additional evidence comes from multiple system atrophy, a parkinsonism-related disorder characterized by DA neuron loss and α-synuclein accumulation in oligodendrocytes, which results in the formation of glial cytoplasmic inclusions^30^. Notably, PD brains show alpha-synuclein inclusions in oligodendrocytes, similar to those found in multiple system atrophy (MSA) brains^31^. While these studies focused on the substantia nigra of PD patients, our research highlights oligodendrocyte changes in the striatum in response to the degeneration of DAergic arbors. These striatal oligodendrocyte changes, which are more specific to our 6-OHDA-induced model, align with prior findings that neurotoxin-induced DA depletion in mice and nonhuman primates induces reactive striatal oligodendrogliosis, which is correlated with the extent of DAergic axon loss^32^. As the roles of astrocytes and oligodendrocytes in PD have gained increasing recognition^33^, our study provides a resource that might be useful for understanding PD-related effects on striatal glial populations.

DA neuron subtypes exhibit differential vulnerability in PD, with DA neurons projecting to the DS being more vulnerable than those projecting to the VS^34,35^. However, it remains unclear whether the transcriptomic changes observed in SPNs are solely explained by the magnitude of DA neuron loss along the dorsoventral gradient of the striatum. In our study, we used spatially defined markers and ST analysis, which revealed that DS clusters, particularly the DMS cluster, are more vulnerable to DA depletion. However, this vulnerability profile did not align with the DAergic axon degeneration pattern. The PD models we used in our study exhibited distinct degeneration patterns, with TH loss in the striatum seemingly dependent on the specific disease model. These variations are mirrored in the transcriptome profiles, likely highlighting distinct underlying pathophysiological mechanisms associated with the mild 6-OHDA and MitoPark models. Despite this variability, the DMS consistently emerged as the most vulnerable spatial cluster, whereas ventral regions such as the ACB and SAMY were the least affected. These findings indicate that the inherent properties of striatal subregions influence their response to DA depletion, which is not entirely dependent on the magnitude of local DA loss. In line with our findings, previous studies have demonstrated differential transcriptional responses to total DA depletion induced by 6-OHDA among striatal regions, as well as in combination with levodopa (L-DOPA) treatment^36–38^. Similarly, spatially distinct responses to DA-releasing agents, such as psychostimulants, further support the intrinsic ability of striatal subregions to adapt to DA content^39–42^. However, in most prior studies, visual parcellation of striatal subregions was used rather than the unbiased molecular parcellation approach we used in our study^36–38,40^. Thus, we provide a detailed molecular resource for understanding the spatial vulnerability of SPNs, offering novel insights into region-specific responses to DA depletion in PD.

The pathophysiology of PD clinical features is typically explained by the distinct functions of the dSPN and iSPN pathways^43–45^. A key finding of our study is the identification of SPN subtypes with distinct vulnerabilities and resilience to DA depletion. We observed that SPNs within the dorsal striatal matrix compartment are the most vulnerable, whereas ventral striatal clusters exhibit the greatest resilience to DA degeneration. Interestingly, one of the most resilient subtypes we identified is the newly characterized eSPN type, despite its localization in the DS. Notably, we found that the equivalent human eSPNs presented minimal transcriptional changes in PD striata, which corresponds to early disease stages in mild 6-OHDA lesion models and middle-stage MitoPark mice. Nevertheless, our pseudotime and trajectory analysis^14,15^ in SPN clusters revealed opposing results between humans and mice. In the human dataset, eSPNs appeared as the endpoint of a lineage that followed Braak stage progression, whereas in the mouse dataset, all lineages originated from early pseudotime points corresponding to MitoPark age. Higher pseudotime values suggest that a cell type has reached a relatively stable transcriptional state under specific conditions. Given that our human samples are likely to represent a more advanced stage of degeneration compared to middle-stage MitoPark mice, this species discrepancy in eSPN pseudotime values may reflect a key difference in disease progression. Specifically, eSPNs may remain relatively unaffected until DAergic degeneration reaches a terminal stage, explaining their later involvement in the human dataset. Evidence supporting the relative resilience of eSPNs to DA loss comes from a study showing that Drd1+/ Drd2+ co-expressing SPNs have dendritic arbors that are less affected by severe DA deafferentation compared to dSPNs and iSPNs^46^. The role of Drd1+/Drd2+ co-expressing SPNs in behavior has recently gained attention^47^. A new study characterized their distinct electrophysiological response to DA^47^, a feature that may underlie their resilience to the DAergic degeneration we observed. Therefore, the contribution to PD symptoms of this recently identified SPN subtype remains to be further explored. We suggest that this cellular resilience profile is a conserved striatal feature in PD in both mice and humans and that eSPNs represent a SPN subtype with significant resilience to the early loss of DAergic innervation across species.

Together, our findings establish a cross-species framework for defining the cell-state dynamics in the parkinsonian striatum, and uncover the identity and response of selectively vulnerable and resilient cell types, thereby revealing cellular principles of susceptibility during PD progression. We envision that this resource in combination with other efforts to map cell states and transcriptional changes can inspire the development of new therapeutic strategies aimed at preserving or restoring function in specific striatal cell types affected by Parkinson’s disease.

## Methods

All details of key lab materials used and generated in this study are listed in a Key Resources Table on Zenodo (https://doi.org/10.5281/zenodo.15065318). Protocols associated with this work can be found on protocols (https://www.protocols.io/file-manager/4B22DCEE066911F0A9360A58A9FEAC02).

### Animals

All details of key lab materials used and generated in this study are listed in a Key Resources Table on Zenodo (https://doi.org/10.5281/zenodo.10553375). Protocols associated with this work can be found on protocols io (https://doi.org/10.17504/protocols.io.dm6gp3k25vzp/v1). All procedures and experiments on animals were performed according to the guidelines of the Stockholm Municipal Committee for animal experiments and the Karolinska Institutet in Sweden (approval numbers N166/15, 155440-2020 and 7362-2019). Adult wild-type mice (C57BL/6J; Charles River) mice aged 3-5 months were used for 6OHDA injections and sacrificed two weeks later, while MitoPark (genotype +/*DAT-cre, TfamloxP/TfamloxP*,^12^) mice were used as follows: Wild-type mice were aged matched with Knock Out animals, different timepoints were used: 11 weeks old, 15 and 18 weeks old. Animals were group housed, up to five per cage, in a temperature (23 °C) and humidity (55%) controlled environment in standard cages on a 12:12 h light/dark cycle with ad libitum access to food and water. The exact number of animals (N) for each experiment is reported in the corresponding figure legend.

### Human Tissue

Fresh-frozen post-mortem brain tissue was acquired from the Cambridge Brain Bank (with the approval of the London—Bloomsbury Research Ethics Committee; 16/LO/0508). Brain slices from the right hemisphere were flash-frozen and stored at −80 °C.

### Stereotaxic injections

All stereotaxic injections were performed on 3-to 5-month-old wild-type C57BL/6J mice. Mice were pretreated with desipramine (25 mg/kg, i.p.; Cat# D3900, Sigma–Aldrich) and pargyline (5 mg/kg, i.p.; Cat# P8013, Sigma–Aldrich) before being placed in a stereotaxic frame, they were anesthetized with 5% isoflurane and secured in a stereotaxic apparatus (Harvard Apparatus). A glass micropipette connected to a Quintessential Stereotaxic Injector (Stoelting) was used to deliver 1 μl of 0.6 mg/ml 6-OHDA dissolved in 0.2% ascorbate, (Cat# 162957, Sigma–Aldrich) into the right medial forebrain bundle (coordinates: AP: -1.2 mm, ML: -1.25 mm, DV: -4.75 mm from the dura). Vehicle solution (0.2% ascorbic acid) was injected as a control. Post-operative care included daily monitoring for one week, with sweetened milk (1:3 dilution in tap water) provided in the home cage to support recovery. Mice were killed two weeks post-surgery for brain collection for cryosectioning or Striatum extraction. Brain tissues were harvested and placed into 2 ml Eppendorf tubes for snap freezing in dry ice, then stored in −80 °C freezer for sample processing.

### Immunofluorescence

Sections were washed once in PFA 4% and then incubated in blocking buffer (10% donkey serum, 0.3% Triton-X in PBS) for 1 hour. Following this blocking step, the sections were incubated for 1 hour at RT with primary antibodies: rabbit anti-TH (Cat# ab152, Abcam) diluted in 2.5% donkey serum and 0.3% Triton-X in PBS. Lastly, sections were washed three times for 5 minutes each in PBS and incubated with fluorescent-conjugated secondary antibodies (Cy-3 anti-rabbit, Jackson Immuno Research, Cat# 711-165-152) for 1 hour. Following three additional 5-minute washes in PBS, sections were mounted on coated slides and covered with coverslips using Mowiol mounting medium. Imaging was performed using a Leica DM6000 B microscope (Leica Microsystems) with a 10× objective.

### Spatial Gene Expression

#### Sample preparation

Six brains in total, three from each group (6OHDA injected mice and vehicle injected mice), were used for Spatial Transcriptomics in this study. The injected hemisphere of each of the brain samples was further used for the Visium experiments. Each sample hemisphere was embedded in O.C.T. and cryo-sectioned in the coordinates corresponding to the interested area (four section per brain were collected at four different bregma coordinates) at 10 µm thickness. The sections were then randomly placed onto Visium Gene Expression arrays for downstream analysis and kept at -80°C for less than a week.

#### Visium Spatial Gene Expression technology and sequencing

The tissue sections mounted on Visium slides were fixed in pre-chilled methanol at -20°C for 30 minutes, in accordance with the 10X Genomics protocol [CG000160 Rev B]. Following fixation, hematoxylin was applied for 5 minutes, followed by a 45-second application of eosin to enhance contrast. Brightfield images were captured using a Zeiss Axiolmager.Z2 microscope equipped with Metafer 5 software. We optimized the tissue permeabilization process specifically for the striatal region, following the 10X Genomics tissue optimization guidelines [CG000238 Rev D]. Preliminary testing identified 14 minutes as the optimal permeabilization time for this brain region. Spatial gene expression profiling was conducted using the Visium platform, strictly following the manufacturer’s instructions [CG000239 Rev D], generating a total of 24 libraries. Sequencing of the 24 libraries was performed on a NextSeq2000 platform, with cDNA libraries diluted to a final concentration of 2 nM and pooled for sequencing. Each sample was sequenced to a depth of 150-200 million reads, with read lengths configured as 28 bp for Read 1, 10 bp for both i7 and i5 indices, and 90 bp for Read 2, in line with the Visium protocol [CG000239 Rev D].

### Chromium single-cell technology and sequencing

#### Nuclei isolation of mouse tissue

Striatum from one hemisphere of 19 brains was collected for nuclei isolation. Brain tissue was placed into a tube together with lysis buffer (10 mM Tris-HCl, 10 nM NaCl, 3 nM MgCl2, 0.1% Nonidet P40 Substitute 0.1%, 1 mM DTT, 1U/µl RNAse inhibitor and Nuclease free water), homogenized by pestle homogenization in a tissue grinder tube and incubated on ice for 5 min. The nuclei were extracted by following 10x Genomics protocol for Nuclei Isolation from Complex Tissues for Single Cell Multiome ATAC + Gene Expression Sequencing (User Guide, CG000375 Rev A). Each sample was filtered through a 70 µm cell strainer, sorted and permeabilized nuclei were pelleted at 500 × g for 5 min at 4 °C, and resuspended in 1 ml PBS 1x, 1% BSA + 1 U/μl RNase inhibitor, for two times. After the second centrifuge nuclei were stained with 7-AAD Ready Made Solution (SML 633-1ML) 0.01% for FACS sorting.

#### Cell sorting of mouse tissue

The stained cell nuclei suspension samples were then analyzed and sorted utilizing a BD FACSAria Fusion flow cytometer using a 100 μm nozzle. Flow cytometry analyses and sorting were carried out by the following gating strategy: nuclei: singlets:7-AAD positive events. The nuclei sub-population was characterized in the Side Scatter-Forward Scatter plot by back-gating from the 7-AAD (ex. 488 nm, em. 692 nm), singlets were gated in the Side Scatter – Pulse width plot and finally the nuclei were gated in the Side Scatter – 7-AAD plot. The nuclei were collected in a BSA coated Eppendorf tube.

#### Chromium sequencing

The nuclei were processed according to the 10X Genomics Chromium Next GEM Single Cell 3’ Reagent Kits v3.1 (Dual Index) protocol (User Guide CG000315 Rev C). For each sample, we aimed to recover between 2,000 and 7,000 nuclei. Libraries were constructed for sequencing in accordance with the protocol and manufacturer’s recommendations. Sequencing was performed using an Illumina NextSeq 2000, targeting a depth of 300 to 400 million reads per sample. The sequencing read lengths were as follows: Read 1, 28 cycles; i7 and i5 index, 10 cycles each; Read 2, 90 cycles.

#### Nuclei Isolation of human putamen

The nuclei isolation from human putamen tissue was performed as previously described^48,49^ using a sucrose gradient-based method. In brief, the tissue was thawed and dissociated in ice-cold lysis buffer (0.32 M sucrose, 5 mM CaCl2, 3 mM MgAc, 0.1 mM Na2EDTA, 10 mM Tris-HCl, pH 8.0, 1 mM DTT) with a 1 ml tissue douncer (Wheaton). The lysate was then carefully layered on top of a sucrose cushion (1.8 M sucrose, 3 mM MgAc, 10 mM Tris-HCl pH 8.0, and 1 mM DTT diluted in milliQ water) and then centrifuged at 30,000 × g for 2 hours and 15 min. Once the supernatant was removed, the pelleted nuclei were softened for 5-10 min in 50 μl of nuclear storage buffer (15% sucrose, 10 mM Tris-HCl pH 7.2, 70 mM KCl, and 2 mM MgCl2, all diluted in milliQ water), and then resuspended in 300 μl of dilution buffer (10 mM Tris-HCl pH 7.2, 70 mM KCl, and 2 mM MgCl2, diluted in milliQ water). The resuspended nuclei were filtered through a cell strainer (70 μm) and then sorted via FANS (with a FACS Aria III, BD Biosciences) at 4° C at low flow rate using a 100 μm nozzle (reanalysis showed >95% purity).

Detailed protocol can be found at DOI: *dx.doi.org/10.17504/protocols.io.5jyl8j678g2w/v1*.

### Pre-processing and Clustering of Spatial Transcriptomics (ST) and snRNAseq Data

#### Spatial Transcriptomics (ST) data analysis

##### Gene count matrix generation from Visium

The sequenced libraries from the Visium experiments were processed using the 10x Genomics Spaceranger pipeline (10x Genomics Space Ranger v1.3.1) to generate the gene count matrices for the downstream data analysis. For the mapping, the Mus musculus reference genome (mm10-2020-A, prebuilt reference from 10X genomics; GENCODE Release M23; Ensembl version Ensembl98) was used.

##### ST data downstream analysis

The downstream analysis was performed in R (v4.1.3; later updated to v4.3.2) using the Seurat package^50–54^ (v5.0.1). In order to load the count matrices in R, Seurat’s Load10X_Spatial() was used. Corresponding samples’ metadata was extracted from previously prepared sheets (see Data availability) and added to the seurat object using the Seurar’s AddMetaData() function. In order to inspect the quality of the dataset, the genecount per spot and UMI per spot distribution was plotted for all samples (i.e., brain sections) using SpatialFeaturePlot(). The quality of the dataset and individual samples was further evaluated using violin plots generated using VlnPlot() from Seurat. In the filtering step, mitochondrial genes, ribosomal genes and hemoglobin genes were filtered out from the samples. Additionally, spots with gene count less than 200, and UMI count less than 100 were also excluded from further analysis. After filtering, to normalize the samples across the dataset, SCTransform() was applied on the ‘Spatial’ assay with additional parameters (glmGamPoi as method, return.only.var.genes set to False, and var.to.regress set to genecount). Top 5000 integration features were used in the integration step and these features were later used to identify integration anchors in the dataset. These integration anchors were the main input to integrate the dataset before running dimensionality reduction. Principal Component Analysis (PCA) method was applied to the integrated dataset using RunPCA() that is provided as a built-in function in the Seurat package. ElbowPlot() was used to inspect the 50 principal components (PCs) in order to select a subset (35 PCs) for the rest of the downstream analysis pipeline. The Harmony package was used for batch effects correction on the dataset and was implemented via the RunHarmony() function. Variations from different samples originating from different Visium capture areas were the major source of batch effects in the dataset which were corrected using Harmony. Clustering of the spots was done using FindNeighbors() and FindClusters() with additional parameters (dims in FindNeighbours() and pc.use in FindClusters() set to 35 PCs, resolution set to 1.5) which resulted in a total of 32 spatial clusters across the whole dataset. VlnPlot(), SpatialDimPlot() and DimPlot() functions were used to visualize the distribution of the spatial clusters in the dataset as well as overlayed on the brain sections. FindAllMarkers() was used on the SCT assay with default parameter settings to identify the markers for each spatial cluster which were later used to assign cluster identities.

##### Subclustering of the striatal clusters in the ST data

First, the spatial barcodes of the spots that belonged to the striatal cluster identity were extracted. These barcodes were then used to subset the seurat object that was originally processed and filtered as per the criteria mentioned in the section ‘ST data downstream analysis’ above. This subset which now represents the dataset belonging to the striatal identity was then normalized using SCTransform with the same settings as those applied to the main ST dataset. Integration of the subset was done in the same manner as the main ST dataset. 40 principal components (PCs) were used in the clustering of the subset (resolution set to 0.7) which resulted in a total of 14 spatial subclusters within the striatal identity. Just like in the main ST dataset clustering, FindAllMarkers() was used to identify the markers for each subcluster using default parameter settings.

##### Differential expression analysis for the ST data clusters

To identify the differentially expressed genes (DEGs) between the vehicle injected and the 6OHDA injected sections for each spatial cluster, FindMarkers() was used (min.pct set to 0.1, min.cells.group set to 3, recorrect_umi set to FALSE). An additional adjusted p-value filter of 0.0001 was later applied to get a significant set of DEGs per cluster for 6OHDA injected versus vehicle mice sections. In the case of DEGs for the striatal sub-clusters, the same functions and parameters were applied but with slight modifications (min.pct set to 0.05).

### Single nuclei sequencing data generation and analysis

#### snRNA sequencing data analysis

We generated the count matrix from the sequencing data for each sample (fastq) using the software commercialized by 10x Genomics Cellranger (10x Genomics Cell Ranger v7.1.0)^55^. Data analysis for the snRNA-seq data was performed in R (v4.3.2) using the packages Seurat^50–53^ (v4.1.0). In total, extracted single nuclei from 20 brain hemispheres from mouse brain were and 54 from human postmortem putamen were processed, resulting in 166810 nuclei before filtering in the mouse dataset and 113650 for the human and used in the analysis, the analysis was performed separately but using the same pipeline. The snRNA-seq assay was generated using Read10X_h5() available in the Seurat package. Malat1, ribosomal,mitochondrial and X and Y chromosome related genes were removed, and the count matrix was further filtered on the number of reads, genes and peaks. Filtering for low quality nuclei and genes was applied separately for each sample. Doublet finder was used for doublet removal (v2.0.3) with a doublet score cut-off of 0.6. RNA-seq data was normalized using SCTransform with V2 regularization, while also regressing out cell cycle effects using cell cycle genes list available in the Seurat package. Harmony (group.by.vars=”Sample” and “Sample_group”) was then used to remove batch effects and integrate data from different samples. Clustering of the spots was done using FindNeighbors() and FindClusters() with additional parameters (dims in FindNeighbours() and pc.use in FindClusters() set to 42 PCs, resolution set to 1.5 for the mouse dataset, and 42 PCs, resolution set to 1 for the human dataset). In the mouse dataset 22 clusters and in the human 30 clusters were found with FindAllMarkers() which was used on the SCT assay with default parameter settings to identify the markers for each cluster which were later used to assign cluster identities. Marker genes were used to annotate clusters, including known markers for glial cells (Microglia, Astrocytes, Oligodendrocytes, and Oligodendrocyte precursors) and neuronal types (Medium Spiny Neurons, Interneurons, and Cortical Neurons). Human dataset followed the same pipeline and analysis, Harmony was run for the same group.vars, with an increase in the correction for the Sample_group (theta=2 instead of theta =1).

#### Gene and cluster annotation

Established marker genes for each cell type were used to annotate the clusters, we used known marker genes for glia (Microglia, Astrocytes, Oligodendrocytes and Oligendrocytes precursors) and neuronal types such as spiny projection neurons, Interneurons, Cortical neurons. The genes are: Mbp, Tnc, Cx3cr1, Slc1a3, Sox1, Pdgfra, Gfap, Aldh1l1, Slc17a7, Slc17a6, Slc32a1, Gad2, Sst, Npy, Pvalb, Chat, Ache, Slc18a3, Gpr6, Ppp1r1b, Rgs9, Gpr88, Adora2a, Drd2, Penk, Pdyn, Tac1, Drd1, Col11a1, Sema5b, Oprm1, Th. In addition to the manual annotation we used the Allen brain Institute tool, MapmyCells^9^ that allowed us to map the cells to their database. The clusters were subsequently annotated and differential expression analysis between clusters was rerun.

#### Subclustering

Data from nuclei belonging to SPNs was normalized using SCTransform with the same settings as those applied to the main clusters’ snRNA seq dataset. Integration of the subset was done in the same manner as the main dataset. In the mouse dataset we selected 41 principal components (PCs) that were used in the clustering of the subset (resolution set to 1.5) which resulted in a total of 13 subclusters within the SPNs identity. Just like in the main dataset clustering, FindAllMarkers() was used to identify the markers for each subcluster using default parameter settings, and annotation of clusters was performed with known marker genes for the SPNs and with the MapmyCell tool used above. The human dataset followed the same pipeline and rationale, 42 PCs and a resolution of 1 were set, in addition a threshold to only include samples with more than 30 cells was applied, resulting in 28 samples (18 PD donors and 10 Non PD donors) included in the analysis. Human dataset followed the same pipeline and analysis, Harmony was run for the same group.vars, with an increase in the correction for the Sample_group (theta=2 instead of theta =1). Annotation of the SPNs cell type was performed as above, using known gene for SPNs subtypes (such as Crym, Gpr155, Wfs1, Dlk1, Drd1, Drd2, Adora2a, Tac1, Id4, Epha4, Kremen1, Sema5b, Lypd1, Oprm1, Htr7, Col11a1, Otof, Pcdh8, Col11a1, Otof, Pcdh8, Nxph4, Ryr1, Rreb1, Ntng2, Negr1, Stac, Crym, Gpr155, Dlk1) and with the the Allen brain Institute tool, MapmyCells^9^ that allowed us to map the cells to their database.

#### Differential expression analysis

Differential gene expression between the treated and control conditions in the mouse model (i.e., 6OHDA treated mice vs control mice, MitoPark KO vs control mice, and between PD and healthy donors in the human samples) both in the analysis of the full dataset or when split between Braak stages (4,5 and 6) was performed using MAST, with a mixed model, using sample as random effect. The analysis was done on each cell type group separately, including genes with expression of at least 5% in each comparison group (i.e., treated and control cells in a given cell type group). Genes with FDR adjusted p-value < 0.0001 were considered differentially expressed. To find DEGs, we used the Seurat function FindMarkers() (min.pct set to 0.1, min.cells.group set to 3, recorrect_umi set t FALSE) and additional filters of adjusted p-value (0.0001) and log2FoldChange (−0.2 and 0.2) which were later applied to get a significant set of DEGs per cluster for mild 6OHDA injected and MitoPark 15/18 versus control samples. In the case of the DEGs for the striatal sub-clusters, the same functions and parameters were applied but with slight modifications (min.pct set to 0.05). To avoid bias from unequal cell counts, we randomly downsampled the larger-n condition within each cell cluster to match the smaller one before performing DE analysis. This ensures each comparison uses the same number of cells per condition, standardizing statistical power across comparisons. Additionally, to account for inherent differences in transcriptome complexity between cell types (e.g., glia vs. neurons), we normalized the number of DE genes by the total gene count within each cluster - thereby adjusting for differences in detection sensitivity and baseline expression landscapes.

#### Trajectory inference and pseudotime analysis

To identify genes with expression changes across pseudotime trajectories, we used tradeSeq, an algorithm optimized for trajectory differential expression analysis. This method fits gene expression data along pseudotime using generalized additive models (GAMs), smoothing expression across the trajectory. By testing for statistically significant changes in coefficients along these trajectories, tradeSeq enables detection of genes that vary in expression over time or between branches. Cells were sampled to ensure balanced representation across stages, and variable genes were selected based on biological variance, by splitting the dataset based on Braak stages, ensuring balanced cell sampling across these stages by randomly selecting cells to match the smallest cluster size (313 cells per cluster). For the Gene Selection, we identified variable genes using scran modelGeneVar, selecting the top 3000 genes with the highest biological variance. This selection focused our analysis on genes most likely to vary along the trajectory. Model Fitting with GAMs: Using tradeSeq’s fitGAM function, we fitted generalized additive models to the expression data of these selected genes across pseudotime, while also accounting for cell weights. The GAMs smooth the gene expression, allowing us to model gradual expression changes along the pseudotime. We set 10 knots to capture nuanced expression patterns across the trajectory. Parallel Processing: For efficiency, we enabled parallel processing with BiocParallel::SnowParam, which helped accelerate computations across this large dataset. The result was a sceGAM object containing fitted models, saved for downstream analysis. This model allows us to identify genes with expression patterns that correlate with pseudotime, helping to uncover the key genes driving progression across disease stages.

## RESOURCE AVAILABILITY

### Lead contact

Further information and requests for resources and reagents should be directed to and will be fulfilled by the lead contact, Konstantinos Meletis.

## MATERIAL AVAILABILITY

### Code Availability

All code used and generated in this study is available at GitHub (https://github.com/craggASAP/Striatum_atlas_earlyPD.git) and Zenodo (https://doi.org/10.5281/zenodo.15065317).

### Data Availability

The raw mouse sequencing data (ST and snRNA-seq) and the raw human postmortem sequencing data are available at the ASAP CRN Cloud (Mouse: https://cloud.parkinsonsroadmap.org/collections/single-nucleus-rna-sequencing-of-the-striatum-of-two-murine-parkinsons-disease-models/overview; Human: https://cloud.parkinsonsroadmap.org/collections/jakobsson-20/overview). Detailed information on which raw mouse or human sequencing data and associated scripts were used in each figure can be found at Zenodo (https://doi.org/10.5281/zenodo.15065317).

These materials are released the terms of the Creative Commons Attribution 4.0 International (CC BY 4.0) license

## Author Contributions

Methodology, M.G., I.M., Y.M. and S.F.; Validation: M.G. and I.M.; Software, M.G and Y.M; Formal Analysis, M.G and Y.M.; Investigation, M.G., I.M., Y.M., S.F, S.G., and K.M; Visualization, M.G., I.M. and Y.M.; Resources, M.G, I.M., Y.M., R.G.,J.J., S.G and K.M.; Data Curation M.G, Y.M and I.M.; Writing – Original Draft, I.M. and M.G.; Writing – Review & Editing, I.M. and K.M.; Funding Acquisition, K.M.; Supervision, I.M., K.M., and S.G; Conceptualization, K.M.; Project Administration K.M.

## DECLARATION OF GENERATIVE AI AND AI-ASSISTED TECHNOLOGIES

During the preparation of the text, the authors used ChatGPT-4.0 for language and grammar corrections and they used it as support in the analysis as a tool for code debugging. After using this tool or service, the author(s) reviewed and edited the content as needed and take(s) full responsibility for the content of the publication.

## Supplementary Figures legends

**Figure S1.**
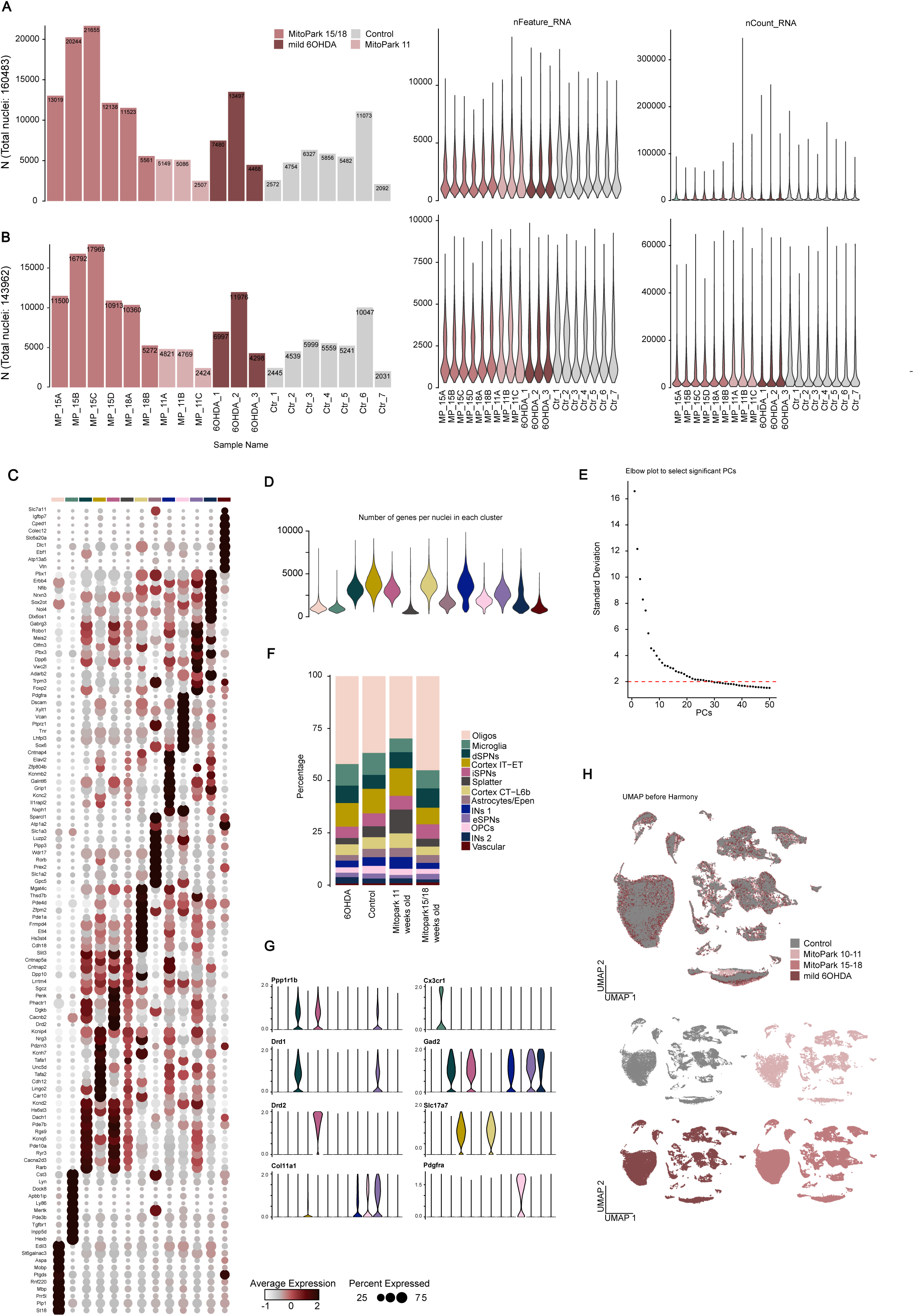
Quality control, clustering, and batch correction of mouse single-nucleus RNA-seq data. **A.** Bar plots of nuclei per sample (left) and violin plots of detected genes and UMIs per nucleus (right), before filtering, color coded by sample group. **B.** Bar plots of nuclei per sample (left) and violin plots of detected genes and UMIs per nucleus (right), after filtering, color coded by sample group (see Methods for criteria). **C.** Dot plots of average normalized expression for the top 10 genes per cluster, colored as in Fig. 1. **D.** Violin plots of the number of detected genes per nucleus across clusters, colored as in Fig. 1. **E.** Elbow plot of the first 50 principal components (PCs), showing explained variance; selected PCs are highlighted in scarlet. **F.** Stacked bar plots of the proportion of nuclei per cluster, split by sample group and colored by cluster as in Fig. 1. **G.** Violin plots of representative marker genes, showing average expression per cluster. **H.** UMAPs before batch correction with Harmony, split and colored by sample group.

**Figure S2.**
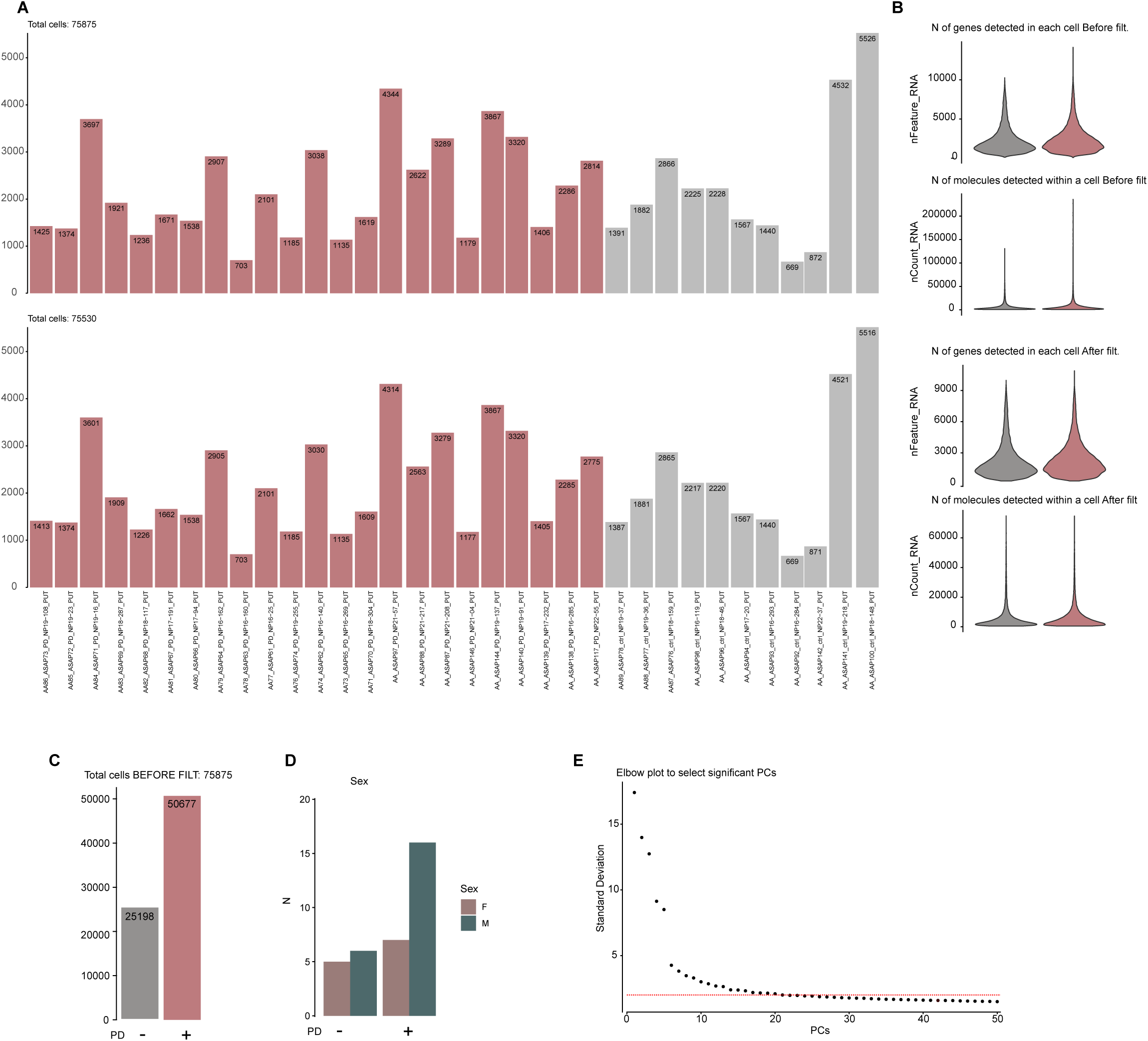
Quality control and filtering of human snRNA-seq data. **A.** Bar plots of nuclei per sample before (top) and after (bottom) filtering, color coded by sample group. **B.** Violin plots of detected genes and UMIs per nucleus, shown before (top) and after (bottom) filtering, color coded by sample group (see Methods for criteria). **C-D.** Bar plots of the total number of nuclei per sample group split by condition (C) and sex distribution across sample groups (D). **E**. Elbow plot of the first 50 principal components, showing explained variance; selected PCs are highlighted in scarlet.

**Figure S3.**
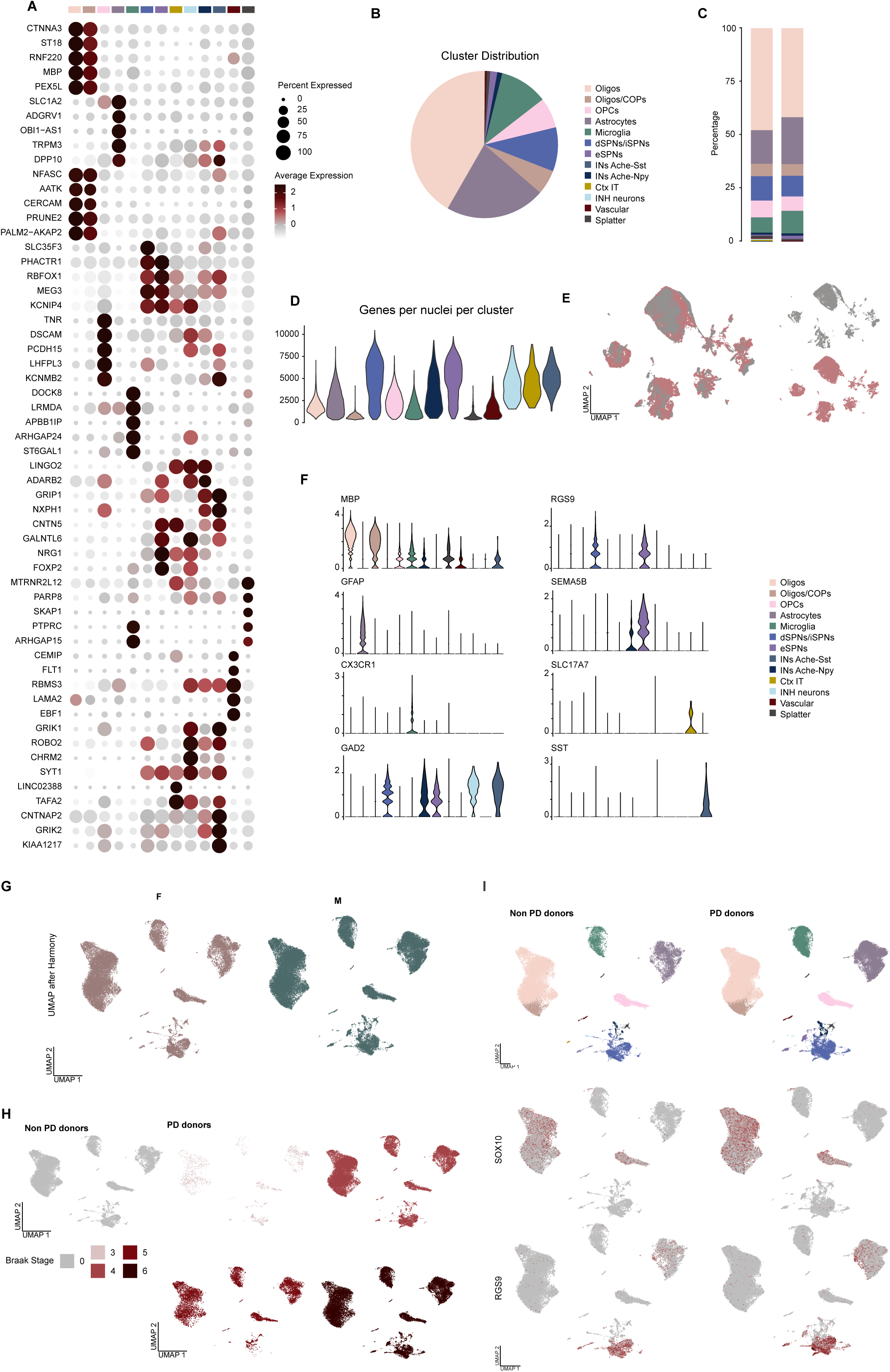
Clustering and batch correction of human snRNA-seq data. **A.** Dotplot showing the average expression level after normalization of top 10 expressed genes per cluster. **B.** Pie chart of nuclei distribution across clusters in the dataset, color coded by cluster as in Fig. 1 **C.** Stacked bar plots of the proportion of nuclei per cluster across sample groups, colored by cluster as in Fig. 1. **D.** Violin plots of the number of detected genes per nucleus across clusters, colored as in Fig. 1. **E.** UMAPs before batch correction with Harmony, colored and split by sample group. **I.** Violin plots of representative marker genes, showing average expression per cluster. **F.** UMAP plot after batch correction, colored by Sex, split by cohort. **G.** UMAPs after batch correction, split by sex. **H.** UMAPs colored and split by Braak stages (non-PD donors labeled as Braak stage 0). **I.** UMAPs split by condition, with clusters and example glial (SOX10+) and stratal marker (RGS9+) genes shown per condition.

**Figure S4.**
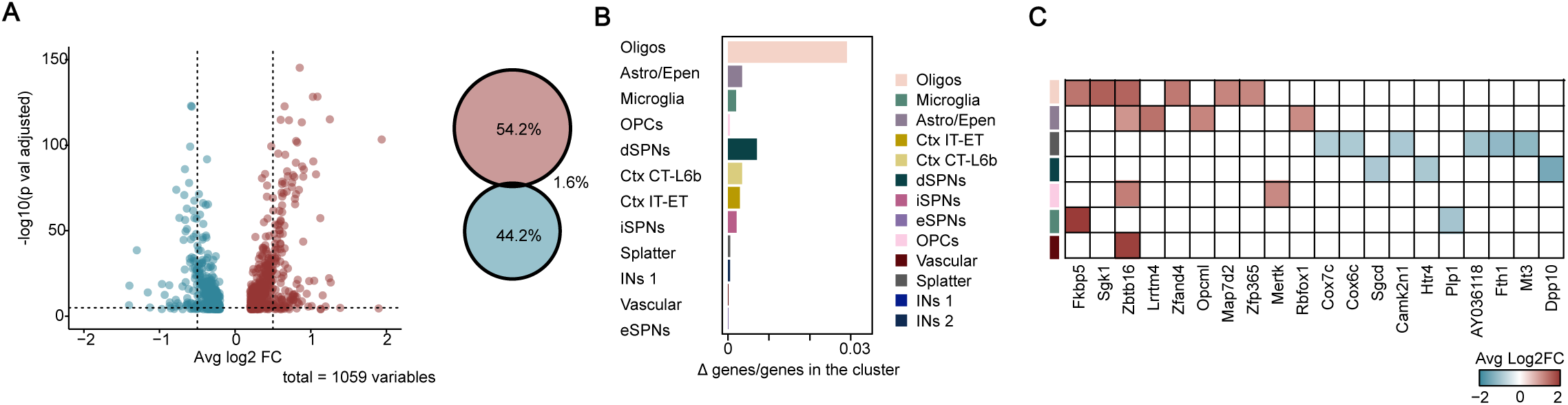
DE analysis and affected striatal clusters in early MitoPark. **A.** Volcano plot of differentially expressed genes in the MitoPark model (10–11 weeks; p < 0.0001, log₂FC > 0.2 or < −0.2, left) and Venn diagram of DE genes (right). **B.** Bar plots of the most affected clusters, ranked by normalized Δ genes/total genes, with neuronal and glial identities indicated. **C.** Heat map of the top 20 dysregulated genes and their cluster assignments, colored by average log₂FC.

**Figure S5.**
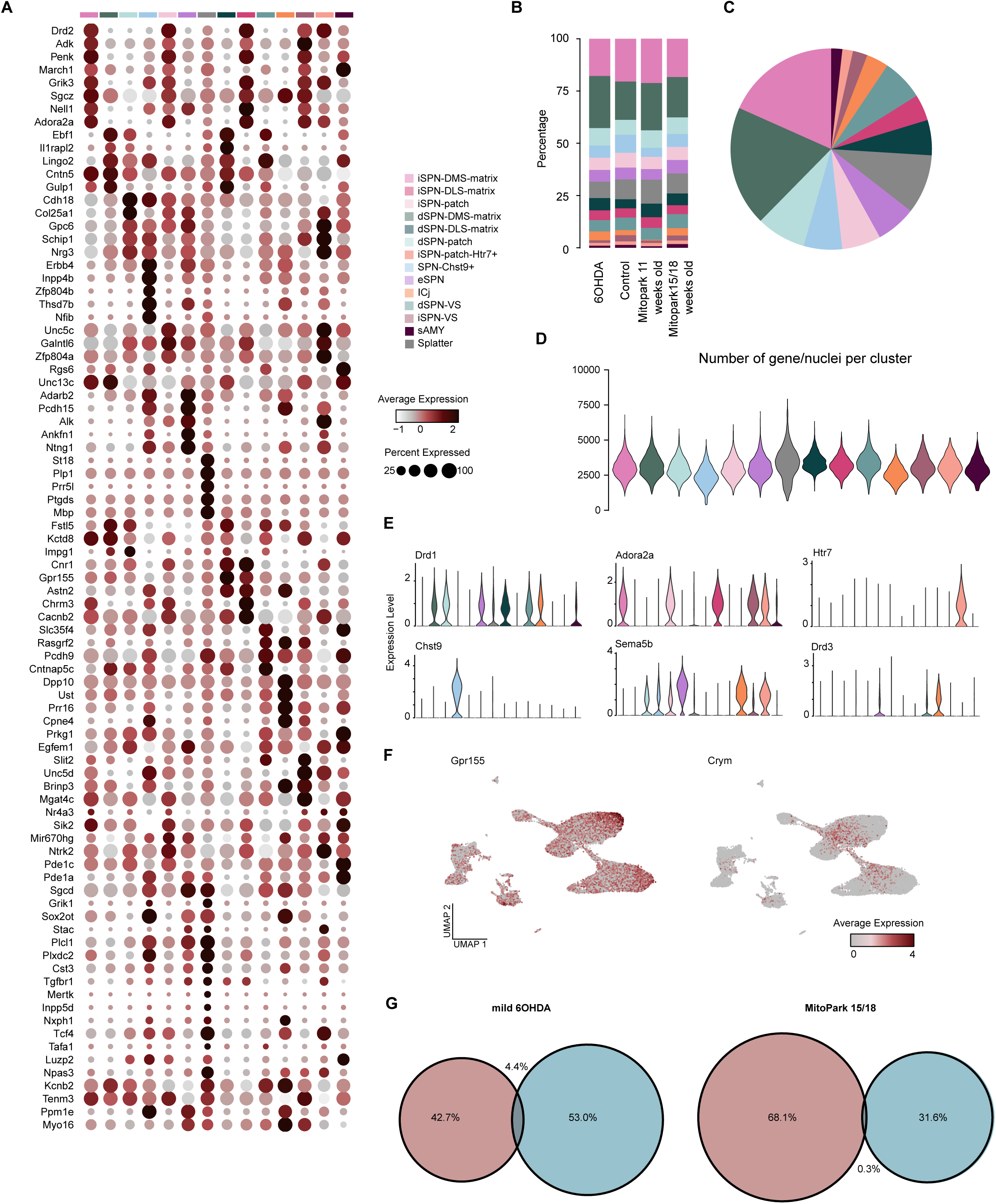
Quality control and marker validation of mouse SPNs subclustering. **A.** Dot plots of average normalized expression for the top 10 genes per cluster in SPNs subclustering. **B.** Stacked bar plots of the proportion of nuclei per cluster across sample groups, colored by cluster as in Fig. 3. **C.** Pie chart of nuclei distribution across clusters in the subclustering dataset, colored by cluster as in Fig. 3. **D.** Violin plots of the number of detected genes per nucleus across clusters, colored by cluster as in Fig. 3. **E.** Violin plots of representative marker genes (one per cluster), showing expression across clusters, colored as in Fig. 3. **F.** Spatial markers of medial (Crym) and lateral (Gpr155) striatum, plotted on UMAPs of SPN subclustering **G.** Venn diagrams of up-and downregulated genes in SPN subclustering, comparing mild 6-OHDA and MitoPark (15–18 weeks).

**Figure S6.**
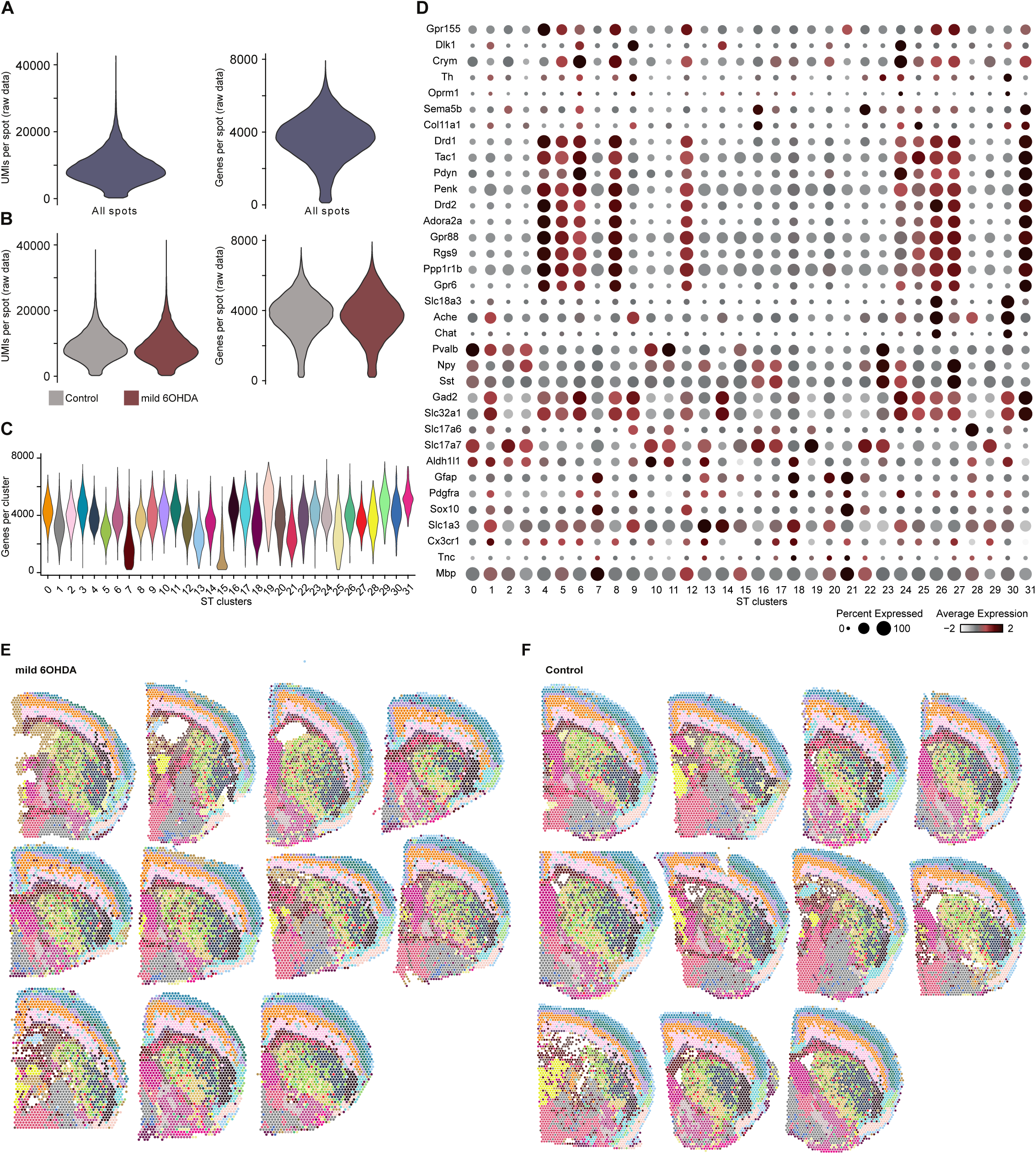
Quality control and clustering of spatial transcriptomic data in mild 6-OHDA model. **A.** Violin plots of detected genes and UMIs across all spots before filtering. **B.** Violin plots of detected genes and UMIs across all spots split by condition (B) (see Methods for filtering criteria). **C.** Violin plots of detected genes and UMIs per sample, colored by sample group, before filtering. **D.** Dot plots of normalized expression for selected marker genes across clusters in the ST dataset. **E–F.** Clusters plotted on spatial spots in control (E) and mild 6-OHDA (F) sections.

**Figure S7.**
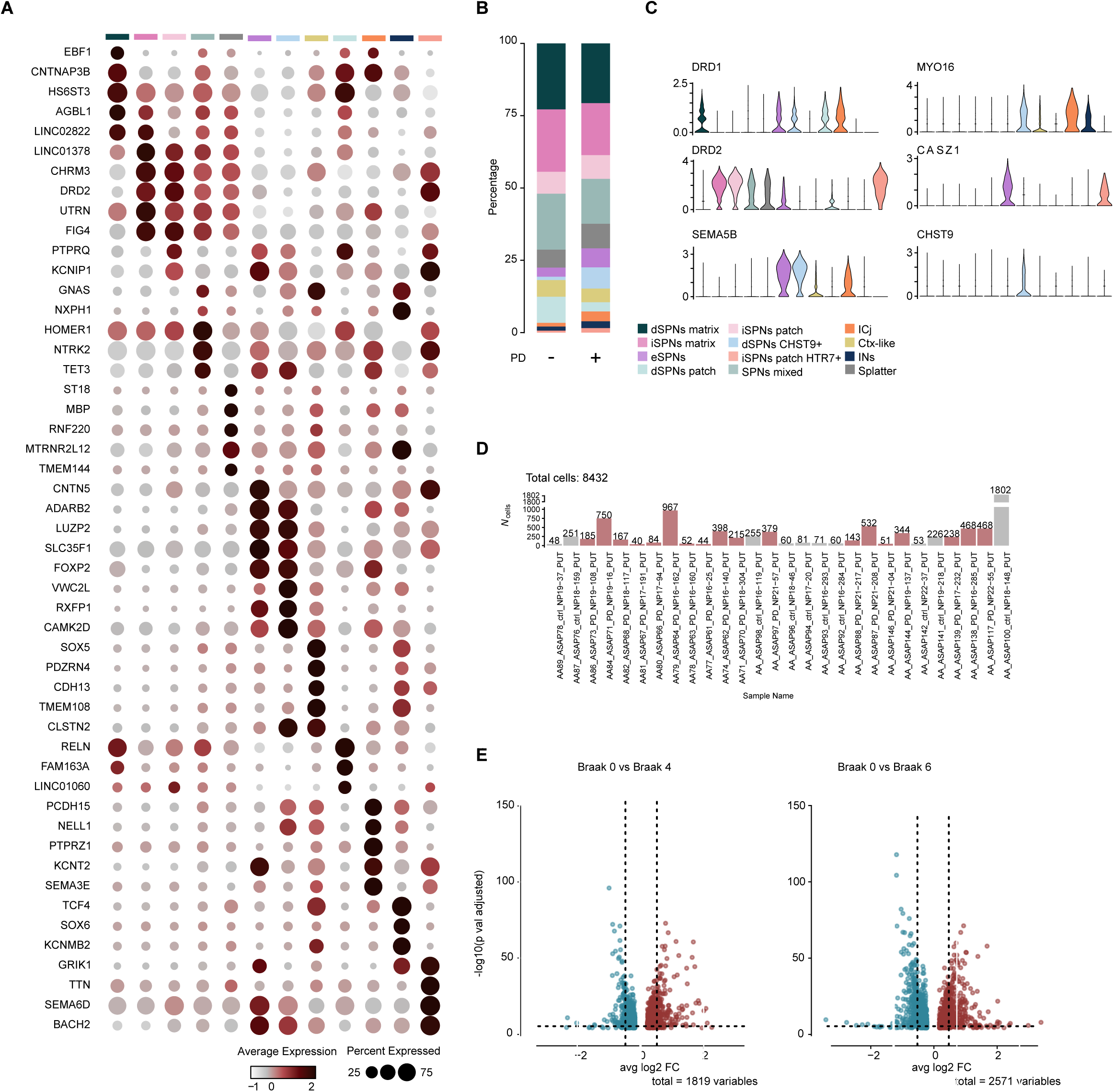
Subclustering and differential expression of human SPNs in PD. **A.** Dot plots of average normalized expression for the top 10 genes per cluster in SPN subclustering. **B.** Stacked bar plots of the proportion of nuclei per cluster across sample groups, colored by cluster as in Fig. 6. **C.** Violin plots of representative marker genes, showing expression across clusters, colored as in Fig. 6. **D.** Bar plots of nuclei per sample after filtering, split and color coded by sample group (criteria in Methods). **E.** Volcano plots of differentially expressed genes in non-PD donors versus Braak stage 4 (left) and Braak stage 6 (right).

